# Using Phylogenomic Data to Explore the Effects of Relaxed Clocks and Calibration Strategies on Divergence Time Estimation: Primates as a Test Case

**DOI:** 10.1101/201327

**Authors:** Mario dos Reis, Gregg F. Gunnell, José Barba-Montoya, Alex Wilkins, Ziheng Yang, Anne D. Yoder

## Abstract

Primates have long been a test case for the development of phylogenetic methods for divergence time estimation. Despite a large number of studies, however, the timing of origination of crown Primates relative to the K-Pg boundary and the timing of diversification of the main crown groups remain controversial. Here we analysed a dataset of 372 taxa (367 Primates and 5 outgroups, 61 thousand base pairs) that includes nine complete primate genomes (3.4 million base pairs). We systematically explore the effect of different interpretations of fossil calibrations and molecular clock models on primate divergence time estimates. We find that even small differences in the construction of fossil calibrations can have a noticeable impact on estimated divergence times, especially for the oldest nodes in the tree. Notably, choice of molecular rate model (auto-correlated or independently distributed rates) has an especially strong effect on estimated times, with the independent rates model producing considerably more ancient estimates for the deeper nodes in the phylogeny. We implement thermodynamic integration, combined with Gaussian quadrature, in the program MCMCTree, and use it to calculate Bayes factors for clock models. Bayesian model selection indicates that the auto-correlated rates model fits the primate data substantially better, and we conclude that time estimates under this model should be preferred. We show that for eight core nodes in the phylogeny, uncertainty in time estimates is close to the theoretical limit imposed by fossil uncertainties. Thus, these estimates are unlikely to be improved by collecting additional molecular sequence data. All analyses place the origin of Primates close to the K-Pg boundary, either in the Cretaceous or straddling the boundary into the Palaeogene.

---

Divergence time estimation is fundamentally important to every field of evolutionary analysis. Reliable estimates of the timing of speciation events across the Tree of Life allow us to correlate these events with both biotic and abiotic phenomena on geological and more recent timescales, thus illuminating those that are most closely associated with periods of rapid diversification (Zhou et al. 2012; Andersen et al. 2015), evolutionary stasis (Alfaro et al. 2009, Liu et al. 2014), or high rates of extinction (Prum et al. 2015; Zhang et al. 2016). Divergence time analysis was first considered feasible with the publication of Zukerkandl and Pauling's (1965) "molecular clock" hypothesis. Very soon thereafter, however, it became clear that there are myriad violations to a uniform clock, and in subsequent decades, increasingly sophisticated models have been developed for relaxing the assumptions of a strict clock (Thorne et al. 1998; Kishino et al. 2001; Thorne, and Kishino 2002; Drummond et al. 2006; Lepage, et al. 2007; Rannala and Yang 2007, Heat et al. 2012). These models can be loosely divided into two categories: autocorrelated models, wherein rates of evolution in daughter species are statistically distributed around the parental rates (Sanderson 1997; Thorne et al. 1998; Aris-Brosou and Yang 2002), and uncorrelated models, wherein each lineage on the tree is free to assume a fully independent rate (Drummond et al. 2006; Rannala and Yang 2007; Paradis 2013).

A parallel challenge to divergence time analysis can be observed in the development of calibration strategies (Marshall 1990; Yang and Rannala 2006; Benton and Donoghue 2007; Marshall 2008; Dornburg et al. 2011). Given that branch lengths on a phylogeny are the product of rate and time, investigators must assume one to infer the other. The most typical method for breaking this impasse is to employ fossil data as calibrations for one or more nodes in a given phylogeny so that the ages of other nodes can be inferred using DNA sequence data. This places an enormous burden on both the correct placement and age assignment of the fossils. If they are misplaced (i.e., assigned to clades where they do not belong) or if their geological ages are misinterpreted, divergence time estimates for the entire tree can be severely compromised (Martin 1993). The uncertainty imposed by paleontological ambiguity has not been as widely appreciated as have been the uncertainties introduced by the finite amount of DNA sequence data, which with the "genomics revolution," is becoming steadily less problematic.

We have reached a state of the art where branch lengths can be estimated with very high precision. The combination of genome-scale data, sophisticated molecular evolutionary models, and ever-increasing computational power has brought us to this point. Advances in DNA sequencing technology are such that virtually every major clade has at least a few species represented by whole-genome sequences, and this trend is rapidly accelerating. Bayesian methods have been developed such that parameter space can be explored during MCMC estimation, and though violations of the molecular clock will continue to present problems, methods for measuring and accommodating rate variation across phylogenies are explicit and generalizable. And finally, the computing power to integrate this information is increasing steadily. But because of the confounding effect of non-independence of rate and time, the expectation of a conventional Bayesian analysis –that infinite data will eventually overcome prior information and provide posterior distributions with certainty – cannot be met (dos Reis and Yang 2013; Zhu et al. 2015).

Thus, the field at present is focused on developing a better understanding of the effects of relaxed clock model choice, and on the impacts of calibration points, both with regard to abundance and placement in the phylogeny. Furthermore, in addressing these challenges, it is an open question as to whether simulation studies or tests of empirical data will be more informative for our understanding of best practices. With regard to clock model choice, an empirical study of three independent datasets showed that autocorrelated models outperform uncorrelated models, though the same study found a "high sensitivity" to the relaxation model employed (Lepage et al. 2007), while another empirical study found, however, that an independent rates model was superior (Linder et al. 2011). Simulation studies have only recently been employed, finding that even with complete taxon sampling, rate autocorrelation is challenging to detect (Ho et al. 2015). This has led to the conclusion that rigorous model selection should be conducted on a case-by-case basis, utilizing a wide range of real data sets, and thus comprising a promising avenue for future research (Ho et al. 2005; Ho and Duchene 2014; Ho et al. 2015).

With regard to fossil calibration strategies, simulation studies (e.g., Duchene et al. 2014) have thus far agreed with previous empirical studies in finding that multiple calibrations are fundamentally important to accurate age estimation (Soltis et al. 2002; Yoder and Yang 2004; Linder et al. 2005; Rutschmann et al. 2007; Marshall 2008). Duchene et al. (2014) noted that calibrations close to the root of the phylogeny are most powerful. They also found that a significant source of error in divergence time estimation relates to misspecification of the clock model, especially when there are few informative calibrations. We cannot stress enough how sensitive posterior time estimates are to fossil information in constructing the time prior (Inoue et al. 2010). For example, different fossil calibration strategies have led to substantially different estimates of divergence times for mammals (dos Reis et al., 2014a) and early animals (dos Reis et al. 2015), regardless of how precise the branch length estimates are (Warnock et al. 2015). Thus, the field has reached the stage wherein there is general agreement that the choice of clock model and calibration strategy are fundamentally important to the accuracy of resulting age estimates, and thus, the way forward will clearly involve both empirical and simulation approaches to the problem.

Here, we hope to contribute to this progress by conducting an exploration of model choice and calibration strategy in a classic empirical system: the Primates. Despite the fact that it is a relatively small and biologically uniform clade, primates have been inordinately and repeatedly the subject of divergence time analysis, with the first studies appearing at the very outset of molecular clock studies (Sarich and Wilson 1967), up to phylogenomic studies encompassing a large set of primate species (Perelman et al. 2011; Springer et al. 2012). This is largely due, undoubtedly, to the fact that we ourselves are members of this clade and can thus be forgiven for a persistent curiosity about our ancestral history. Age estimates for major primate divergence events have varied broadly among different studies (see Table 1), though one result has been relatively constant throughout: primate origins have been typically shown to predate the Cretaceous-Paleogene (K-Pg) mass-extinction event.

**Table 1.** Overview of estimates of divergence times in Primates (in millions of years ago) for selected studies.

| Study | Data/Analysis | Primates | Haplorrhini | Anthropoidea | Platyrrhini | Catarrhini | Homininae | Strepsirrhini | Lorisiformes | Lemuriformes |
| --- | --- | --- | --- | --- | --- | --- | --- | --- | --- | --- |
| Sarich and Wilson (1967) | Immunological distance/strict clock |  |  |  |  |  | 5 |  |  |  |
| Hasegawa, et al. (1985) | 896 bp mtDNA/strict clock |  |  |  |  |  | 3.7 |  |  |  |
| Koop, et al. (1989) | B-globin DNA sequences/strict clock | 55 |  | 40 |  | 25 |  |  |  |  |
| Purvis (1995) | Super Tree Analysis | 57.2 |  | 39.9 |  | 14.7 | 8.1 | 41.8 | 22.1 | 39.6 |
| Kumar and Hedges (1998) | 658 nuclear genes (analyzed individually)/strict clock |  |  |  | 47.6 |  | 6.7 |  |  |  |
| Goodman et al. (1998) | 60-80 Kbp globin gene region/local clocks | 63 | 58 | 40 |  |  | 18 | 50 | 23 | 45 |
| Yang and Yoder (2003) | 2404 bp mtDNA/local clocks |  |  | 57.6 |  |  |  | 69.9 | 38.9 | 64.8 |
| Poux and Douzery (2004) | 1278 bp nDNA/local clocks |  | 56.7, 58.4 |  |  |  |  | 45.4, 46.7 | 13.8, 14.2 | 39.6, 40.7 |
| Eizirik, et al. (2004) | 8,182 bp nDNA/relaxed clock | 77.2 |  | 43.6 |  |  |  | 59.6 |  |  |
| Steiper and Young (2006) | 59.8 Kbp of genomic data/relaxed clock w/ autocorrelated rates | 77.5 |  | 42.9 | 20.8 | 30.5 | 8.6 | 57.1 |  |  |
| Bininda-Emonds, et al. (2007) | 51,089 bp mtDNA & nDNA/ad hoc relaxed clock | 87.7 |  |  |  |  |  |  |  |  |
| Chatterjee, et al. (2009) | 6,138 bp mtDNA & 2,157 bp nDNA/relaxed clock | 63.7 |  | 42.8 | 26.6 | 23.4 | 10.7 | 51.6 | 37.5 | 46.2 |
| Perelman, et al. (2011) | 34,927 bp DNA/relaxed clock w/independent rates | 87.2 | 81.3 | 43.5 | 24.8 | 31.6 | 8.3 | 68.7 | 40.3 | 58.6 |
| Springer, et al. (2012) | 69 nDNA genes; 10 mtDNA genes/relaxed clock w/ autocorrelated rates | 67.8 | 61.2 | 40.6 | 23.3 | 25.1 | 8.0 | 54.2 | 34.7 | 50.0 |
| dos Reis, et al. (2012) | 14,644 genes w/ 20.6 Mpb/relaxed clock w/ autocorrelated rates | 68.2 | 65.0 | 37.4 |  | 26.4 | 10.4 | 55.1 | 35.6 | 49.3 |
| Finstermeier, et al. (2013) | complete mtDNA genomes/relaxed clock w/independent rates | 66.2 | 63.0 | 45.3 | 22.0 | 32.0 | 8.4 | 56.9 | 34.5 | 47.1 |
| Pozzi, et al. (2014) | complete mtDNA genomes/relaxed clock w/ autocorrelated rates | 74.1 | 70.0 | 46.7 | 20.9 | 32.1 | 10.6 | 66.3 | 40.3 | 59.6 |
| This Study (Strategy A) | 3.4 million bp/ relaxed clock w/ autocorrelated rates | 74.4 | 70.6 | 45.0 | 25.3 | 32.6 | 10.5 | 62.7 | 37.9 | 57.2 |
| This Study (Strategy B) | 3.4 million bp/ relaxed clock w/ independent rates | 84.8 | 78.6 | 45.9 | 25.3 | 28.8 | 7.6 | 64.0 | 38.2 | 55.3 |

Notes: All estimates are mean estimates; see original works for confidence/credible intervals; all taxonomic designations signify crown nodes (Primates = the divergence of Strepsirrhini and Haplorrhini; Haplorrhini = the divergence of *Tarsiidae* from Anthropoidea; Platyrrhini; Catarrhini; Homininae = the divergence of *Gorilla* from *Pan+Homo*; Strepsirrhini = the divergence of Lorisiformes and Lemuriformes; Lorisiformes = the divergence of Galagidae and Lorisidae; Lemuriformes = the divergence of *Daubentonia* (the aye-aye) from other Malagasy lemurs.

Our study explores the effects of an autocorrelated versus an uncorrelated rate model on age estimates, and also explores the consequences of two different interpretations of both the age and the placement of key fossils with the living primate radiation. We apply these two strategies to a large phylogenomic dataset for Primates (372 species with 61 thousand base pairs and 9 species with 3.4 million base pairs). Until very recently, reliable calculations of branch lengths and age estimates within an analysis of this magnitude would have been beyond the capacity of computational methods. We have tackled many of these challenges by deploying the sequential Bayesian procedure used by dos Reis et al. (2012) wherein the posterior age estimates derived from a small taxonomic sample with genome-scale data are then deployed as priors for a subsequent analysis with many species and a much-reduced nucleotide sample. This procedure reduces the computational cost of a typical combined data analysis. It also helps to alleviate the concerns with the “missing data” problem, because in the sequential analysis the analysis of times on the genome-scale partition are carried out on a 100% complete data set (although missing data are still present in the second dataset).

The molecular timeline for primate evolution that emerges from this study can be interpreted with confidence. The dataset is sufficiently large to provide highly precise branch length estimates, and the methods used are robust in accommodating violations of the molecular clock. The comparison of the two calibration strategies reveals their impact on the results by giving different age estimates within the tree, though the variation in inferred ages is not extreme. Which ages are considered most accurate will depend in large part on the degree of confidence in the fossils and their placement. As an unanticipated result of the study, the difference in age estimates for the deepest nodes of the phylogeny differ markedly when comparing the molecular rate models, with Bayesian model selection supporting the autocorrelated model. As with previous studies over the past several decades, the ancestral primate lineage is hypothesized to have survived the great K-Pg mass extinction event.

## METHODS

Bayesian estimates of divergence times of Primates were obtained using a supermatrix of molecular data with 372 species and 3.44 million base pairs (Mbp), combined with 17 fossil calibrations. The matrix is the result of merging the 372-species and 61 thousand base-pairs (Kpb) data set of Springer et al. (2012) with a 10-species subset of the genome-scale alignment of dos Reis, et al. (2012). Bayesian analyses were done using the program MCMCTree (Yang 2007). We assessed the robustness of time estimates by varying the clock model (strict clock, independent rates and correlated rates), and by obtaining estimates under two fossil calibration strategies. Note that time estimates were obtained in two steps: In the first step estimates were obtained for the small phylogeny of 10 species with a long alignment (3.38 Mbp). The marginal posterior of times was then used to construct the time prior in the second step for the 372-species phylogeny with a shorter alignment (61 Kbp). This approach is almost the same as analysing the fully concatenated alignment in one step (3.38 Mbp + 0.061 Mbp), but is computationally less demanding. All alignments, tree topology and fossil calibrations are available in Supplementary Material.

### Sequence Alignment and Tree Topology

*Springer et al (2012) alignment.*—We retrieved the sequence alignment of Springer, et al. (2012), which is an extended version of the alignment of Perelman, et al. (2011). The alignment has 372 species (367 primates and 5 outgroup species) and 79 gene segments (69 nuclear and 10 mitochondrial). The composite lagomorph sequence (an outgroup) was removed. We added a scandentian species (*Tupaia belangeri*), because its complete genome is available in the alignment of dos Reis, et al. (2012), and because it has the 10 gene segments from the mitochondrial genome available (accession NC_002521). Many of the nuclear gene segments in the alignment of Springer, et al. (2012) were mixtures of introns, exons and UTRs, with out-of-frame indels in some exons. We manually curated the exons, and separated the coding and non-coding segments of the alignment. These adjustments were necessary to facilitate an informed partition-based analysis of the data. Our modified version of Springer’s alignment was thus divided into 6 partitions: (1) 1st and 2nd codon positions for mitochondrial genes; (2) 3rd positions for mitochondrial genes; (3) mitochondrial RNA genes; (4) 1st and 2nd codon positions for nuclear genes; (5) 3rd positions for nuclear genes; and (6) non-coding segments of nuclear genes (UTRs and introns). The concatenated alignment has 372 species and is 61,132 base pairs long (missing data 27%, Table 2). Our partitioning into codon positions and coding vs. non-coding sequences follows established recommendations (Shapiro et al., 2006; Yang and Rannala, 2006; Nascimento et al., 2017).

**Table 2.** Sequence alignment summary

| **Alignment** | **Partition<sup>a</sup>** | **Sites** | **Species** | **Missing data<sup>b</sup>** |
| --- | --- | --- | --- | --- |
| Springer et al. | 1. mit 1st+2nd | 4,816 | 330 | 61.4% |
|  | 2. mit 3rd | 2,408 | 330 | 61.4% |
|  | 3. mit RNA | 2,169 | 220 | 45.7% |
|  | 4. nuclear 1st+2nd | 16,309 | 239 | 53.8% |
|  | 5. nuclear 3rd | 8,156 | 239 | 53.8% |
|  | 6. nuclear non-coding | 27,274 | 220 | 46.3% |
|  | **Partitions 1–6** | **61,132** | **372** | **51.1% (68.8%)** |
| dos Reis et al. | 7. nuclear 1st+2nd | 2,253,316 | 10 | 0.0% |
|  | 8. nuclear 3rd | 1,126,658 | 10 | 0.0% |
|  | **Partitions 7–8** | **3,379,974** | **10** | **0.0% (97.3%)** |
| **Total** |  | **3,441,106** | **372** | **0.78% (96.8%)** |

a. For topology estimation with RAxML, we used seven partitions: partitions 1 to 3, then 4 and 7 as one partition, 5 and 8 as one partition, and partition 6 divided into two: UTRs and Introns. b. Numbers in brackets are the % missing data for RAxML. Note that MCMCTree only uses the species present in a partition to calculate the likelihood for the partition. In RAxML missing species in a partition are represented as sequences of only gaps in the partition, and thus the amount of missing data is larger.

*dos Reis et al. (2012) alignment.*—We retrieved the genome-scale sequence alignment of dos Reis, et al. (2012) of 36 mammal species, from which we extracted the sequences for 9 primates and 1 scandentian. The dos Reis et al. (2012) alignment was prepared using the highly curated mammalian genomes available in Ensembl. Though three additional primate genomes have become available in this database in the time since the original alignment was prepared, it is unlikely that their inclusion would change our results. The nine species represented in our study provide comprehensive phylogenetic representation of all major nodes in the primate tree and represent each of the higher-level clades (Figure 1, inset). The original alignment has 14,632 nuclear, protein-coding genes, from which we removed 43 genes that were already present in the Springer alignment and 1 gene that was extremely long. All columns in the alignment with ambiguous nucleotides were removed, though care was taken not to disrupt the reading frame of the aligned coding sequences. The alignment was divided into two partitions: (1) 1st and 2nd codon positions; and (2) 3rd codon positions. The final alignment has 10 species and is 3,441,106 base pairs long (missing data 0%, Table 2).

**Figure 1.**
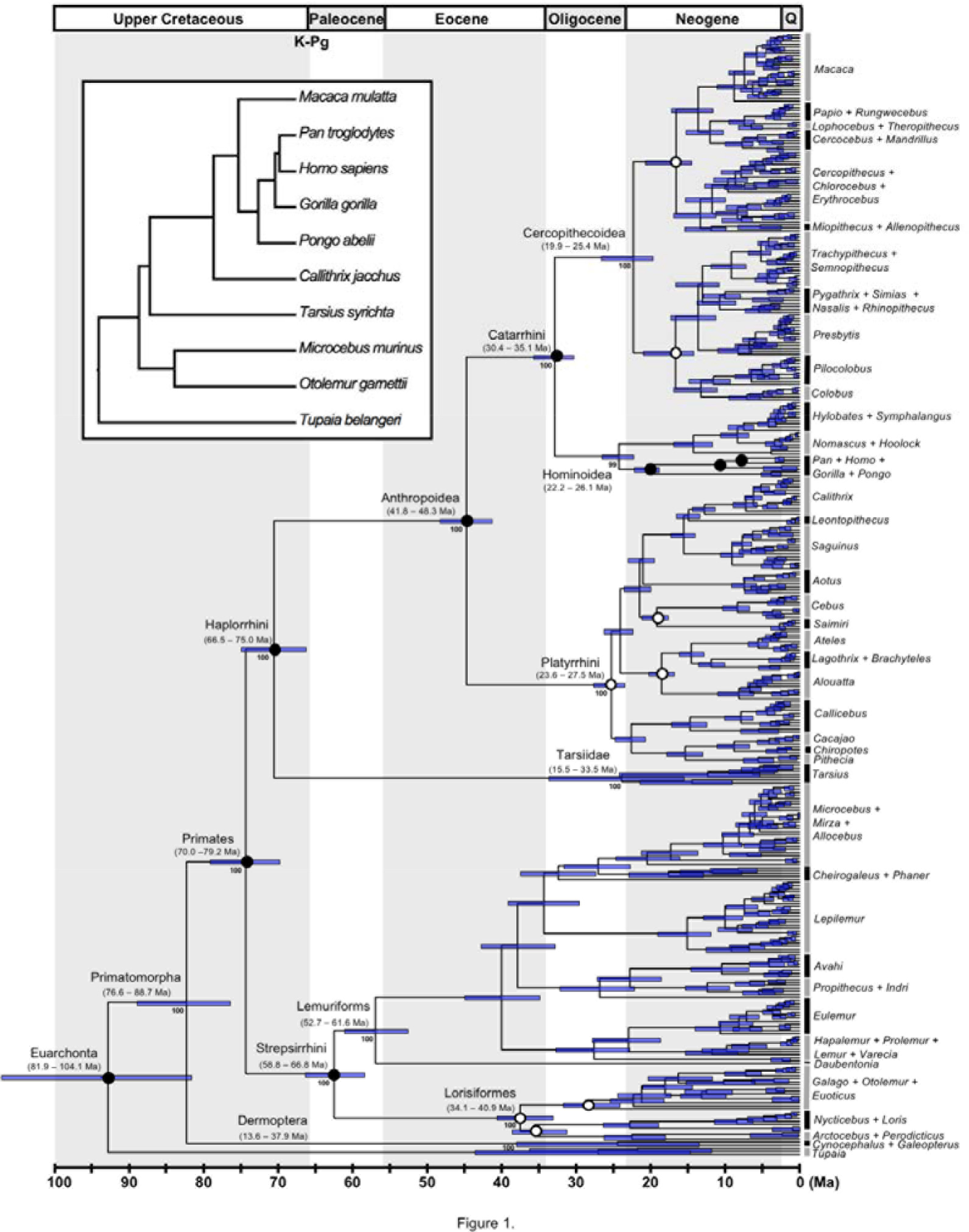
The timetree of Primates. Nodes are drawn at their posterior mean ages in millions of years ago (Ma) estimated under the autocorrelated-rates (AR) clock model and calibration strategy A. Filled dots indicate nodes calibrated with the posterior times from the 10-species tree (inset figure), and empty dots indicate nodes with fossil constraints in the 372-species tree. Horizontal bars and numbers in parenthesis represent the 95% posterior CI for the node ages. Numbers associated with branches are ML Bootstrap support values of major clades.

*Tree topology*.—The topology of the 372-species phylogeny was estimated by maximum likelihood (ML) using RAxML v 8.0.19 (Stamatakis 2014) under the GTR+G model (Yang 1994b, a), using seven partitions (Table 2) and 100 bootstrap replicates.

### Fossil Calibrations and Time Prior

The two fossil calibration strategies used in this study are summarized in Table 3. They represent two different interpretations of the fossil record to construct calibrations for use in molecular clock dating analyses. Calibration strategy A is novel to this study, and calibration strategy B is based on the Primate calibrations of dos Reis, et al. (2012). Detailed justifications for the novel calibrations are provided in Appendix 1.

**Table 3.**
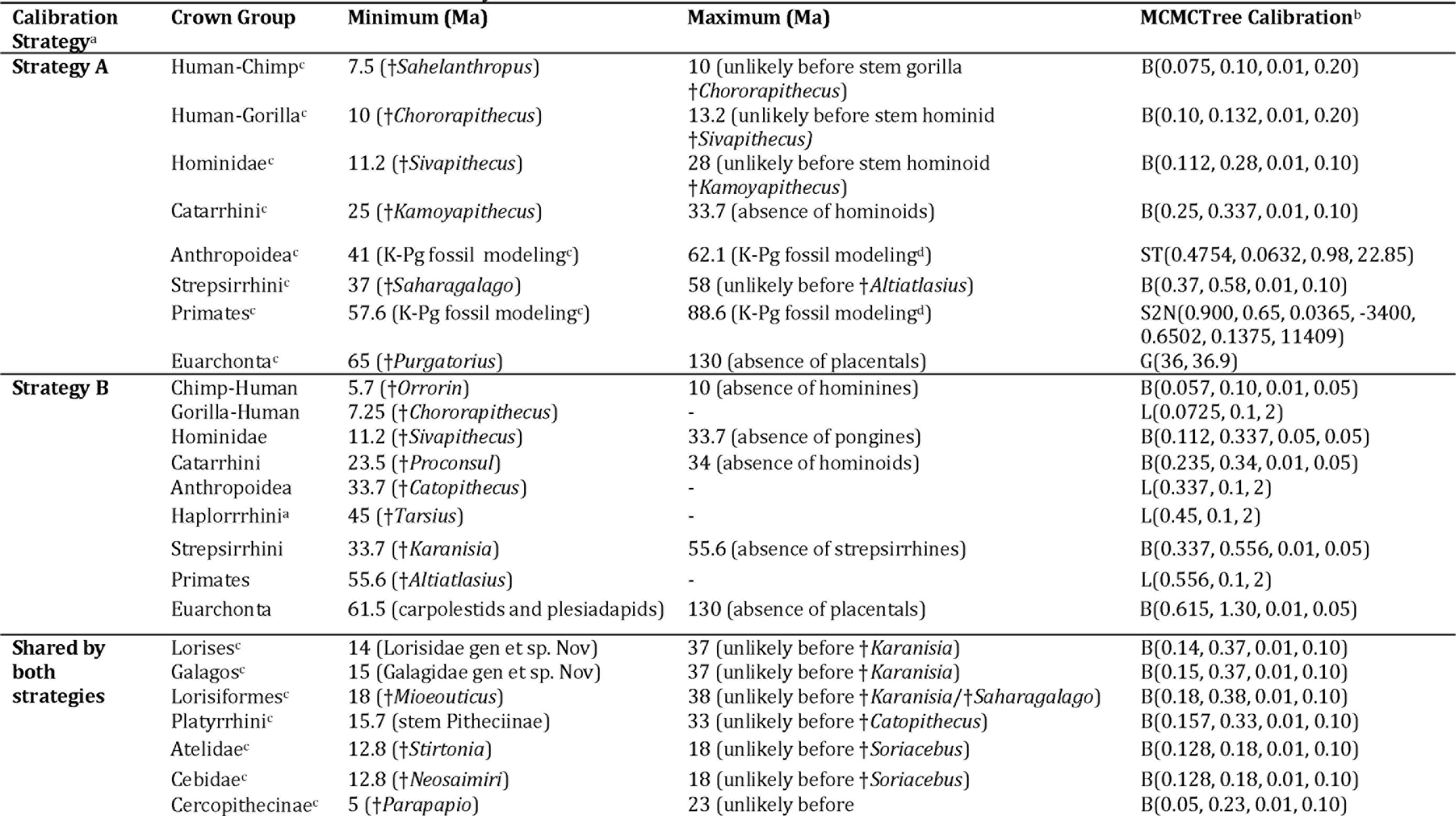
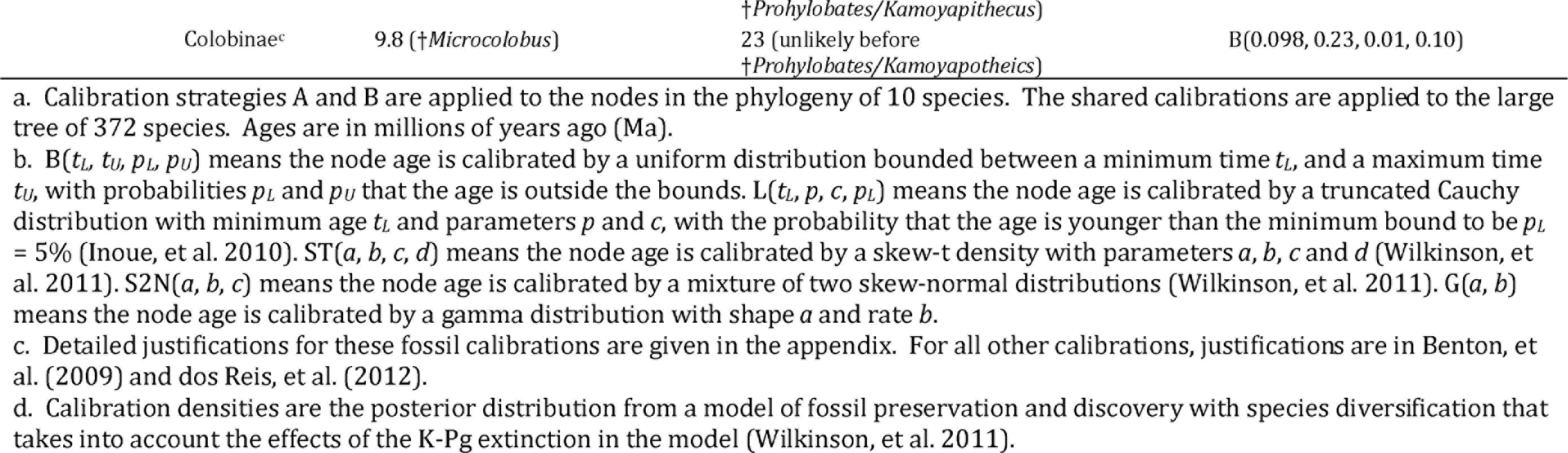
Fossil calibrations used in this study.

*Fossil calibration strategy A*.—We used the fossil-based prior densities constructed by Wilkinson, et al. (2011) to calibrate the ages of crown Primates and crown Anthropoidea. The prior densities were constructed by modelling the processes of speciation, extinction, fossil preservation, and rates of fossil discovery in Primates. The effects of the K-Pg extinction were accounted for in the model. We calibrated six more node ages by using uniform distribution densities with soft bounds (Yang and Rannala 2006). We set the probability of violating a minimum bound to 1%. Because maximum bounds are based on weak evidence, we set the probability that a maximum bound is violated to 10% or 20%. The crown Haplorrhini node was left with no calibration as the branch separating that clade from crown Primates is very short and we wanted to avoid truncation with the fossil-modelling density on crown Primates. The prior on the age of crown Haplorrhini is instead set using the birth-death process with parameters *λ* = *μ* = 1 and *ρ* = 0. These parameter values specify a uniform kernel density (Yang and Rannala 1997, equation 7).

*Fossil calibration strategy B.*—We used the same nine calibrations that dos Reis, et al. (2012) used to calibrate the Primates and Scandentia clades. An additional calibration based on † *Tarsius* sp. was used for the Haplorrhini node. For nodes with a minimum bound only, modelled using a truncated Cauchy density, the spread parameter was set to *c* = 2 (Inoue, et al. 2010). For maximum bounds the probability that the bound was violated was set to 5%. There are other differences between strategies A and B (Table 1). For example, in A, we considered †*Sahelanthropus*, dated to 7.25 million years ago (Ma), to be the oldest member of the human-chimpanzee clade and used it to calibrate the clade accordingly, while in B, dos Reis, et al. (2012) used †*Orrorin* (5.7 Ma) instead. In A, †*Chororapithecus* is given an age of 10 Ma, while in B it is given the younger (perhaps more conservative) age of 7.25 Ma (Benton, et al. 2009). We note that the ages of fossils and their relationships to extant groups are often controversial and cannot overemphasize the degree to which differences of opinion among palaeontologists are an important source of uncertainty in the construction of fossil calibrations, and accordingly, divergence time estimates throughout the phylogeny.

*Calibrating the 372-species phylogeny.*—Strategies A and B were used to obtain time estimates for the 10-species phylogeny using the 3.38 Mbp alignment Then skew-t densities were fitted by ML to the marginal posterior ages of each of the 9 internal nodes in the 10-species phylogeny, and used to calibrate the corresponding nodes in the 372-species tree. Eight additional fossil calibrations (Table 3) were used to calibrate additional nodes in the 372-species tree. For nodes without calibrations, the time prior was constructed using the birth-death process with parameters *λ* = *μ* = 1 and *ρ* = 0. Bayesian time estimation then proceeded on the 372-species tree and 61 Kbp alignment as usual.

### Rate Prior

The time unit is 100 My. For the 10-species analysis, the rate prior was set as follows: the nuclear substitution rate at third codon positions in apes is roughly within 10^−9^ substitutions per site per year (s/s/y) (Burgess and Yang, 2008). At first and second codon positions it is about a fifth of the third position rate, or 2×l0^-10^ s/s/y. This gives roughly an overall rate of about 5×l0^-10^ s/s/y for the three positions combined. We thus used a diffuse gamma density G(2,40) with mean 0.05 and 95% prior credibility interval (CI) 0.00606-0.139 (corresponding to 6.06×l0^-12^ to 1.39×l0^-10^ s/s/y). The analysis was conducted under both the auto-correlated rates (AR) and independent rates (IR) models. Parameter *σ*^2^ in the AR and IR models was assigned a gamma prior G(1, 10). Note that the average rate for loci, *μ*_*i*_, and 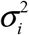 are assigned a gamma-Dirichlet prior (dos Reis, et al. 2014b).

For the 372-species phylogeny, the rate prior was assigned as follows: the mitochondrial substitution rate at third positions is about 20 times the rate at third positions in nuclear genes or 2×l0^-8^. Assuming 1st and 2nd codon positions evolve at about a fifth of the third position rate we get roughly 4×10^−9^. The prior mean is then approximately 2.5×10^−9^ s/s/y, which is the weighted average (by number of sites) of the substitution rates for the nuclear and mitochondrial partitions. We thus used a gamma density G(2, 8) with mean 0.25 and 95% CI 0.0302-0.696. For *σ*^2^ we used G(1, 10). The rate priors are summarised in Table 4.

**Table 4.** Rate priors used in this study. Time unit is 100 million years.

| Alignment | Mean locus rate, $\mu_i$ | $\sigma_i^2$ |
| --- | --- | --- |
| 10-species (nuclear) | G(2, 40); mean $5 \times 10^{-10}$ subs/site/year | G(1, 10) |
| 372-species (nuclear and mitochondrial) | G(2, 8) ; mean $2.5 \times 10^{-9}$ subs/site/year | G(1, 10) |

### MCMC and Bayesian Selection of Clock Model

MCMC analyses were carried out with the program MCMCTree (Yang 2007), using the approximate likelihood method (dos Reis and Yang 2011). Convergence of the MCMC to the posterior distribution was assessed by running the analyses multiple times. MCMCtree runs were carried out without sequence data to calculate the joint prior of node ages. Results from all analyses were summarised as posterior means and 95% CIs.

We have implemented marginal likelihood calculation by thermodynamic integration (path sampling) in the program MCMCTree. This allows us to calculate Bayes factors (BF) and posterior model probabilities to select for a clock model in the analysis. Details
of our implementation are given in Appendix 2. Extensive discussions on marginal likelihood estimation by thermodynamic integration and stepping-stones (a related method) in the phylogenetic context are given in Lartillot and Philippe (2006), Lepage, et al. (2007) and Xie, et al. (2011). A detailed simulation study is given in Ho et al. (2015).

Thermodynamic integration is computationally intensive as we must sample from the power posterior *f*(*θ*) *f*(*D*|*θ*)^*β*^, in a sampling path from the prior (*β* = 0) to the posterior (*β* = 1). Because the approximation to the likelihood is not good when samples are taken far away from the maximum likelihood estimate (dos Reis and Yang 2011), as it happens when *β* is small, the approximation cannot be used in the calculation of the power posterior. Thus, we use exact likelihood calculation on a smaller dataset of nine primate species (inset of Fig. 1), for the six partitions of the Springer alignment (Table 2) to perform the Bayesian selection of clock model. We use 64 *β*-points to construct the sampling path from the prior to the posterior and calculate the marginal likelihoods for the strict clock, and the AR and IR models.

### Effect of genome-scale data

In a conventional statistical inference problem, the variance of an estimate decreases in proportion to 1/*n*, with *n* to be the sample size. Thus, as the sample size approaches infinity, the variance of an estimate approaches zero and the estimate converges to the true value. In divergence time estimation, which is an unconventional estimation problem, the non-identifiability of times and rates means that the uncertainty in the posterior of times does not converge to zero as the amount of molecular data (the sample size) approaches infinity, but rather converge to a limiting value imposed by the uncertainties in the fossil calibrations (Yang and Rannala, 2006; Rannala and Yang, 2007). For infinitely long alignments, an infinite-sites plot (a plot of uncertainty in the time posterior, measured as the width of the CI, i.e., the difference between the 2.5% and 97.5% limits vs. the mean posterior of time) would converge onto a straight line. This line represents the amount of uncertainty in time estimates for every 1 million years (My) of divergence that is due solely to uncertainties in the fossil calibrations. We calculate the infinite-sites plots for time estimates on the 372-species phylogeny to study the effect of genome-scale data on the uncertainty of species-level time estimates.

## RESULTS

### A Timeline of Primate Evolution

The main time estimates for Primates clades are summarised in Table 5, and Figure 1 illustrates time estimates under the AR model and calibration strategy A (Figs. 2 - 4 show detailed timetrees for the major clades). Under calibration strategy A and the AR model, we find that crown Primates originated 79.2-70.0 million years ago (Ma), before the K-Pg event at 66 Ma. However, the diversification of the main clades occurred much later. Crown Anthropoidea originated 48.3-41.8 Ma, with its two main crown groups, Catarrhini (Old World monkeys and apes) and Platyrrhini (New World monkeys) originating at 35.1-30.4 Ma and 27.5-23.6 Ma respectively. Crown tarsiers originated 33.5-15.5 Ma. Crown Strepsirrhini date back to 66.8-58.8 Ma, with its two main crown groups, Lemuriformes and Lorisiformes, dating back to 61.6-52.7 Ma and 40.9-34.1 Ma respectively.

**Figure 2.**
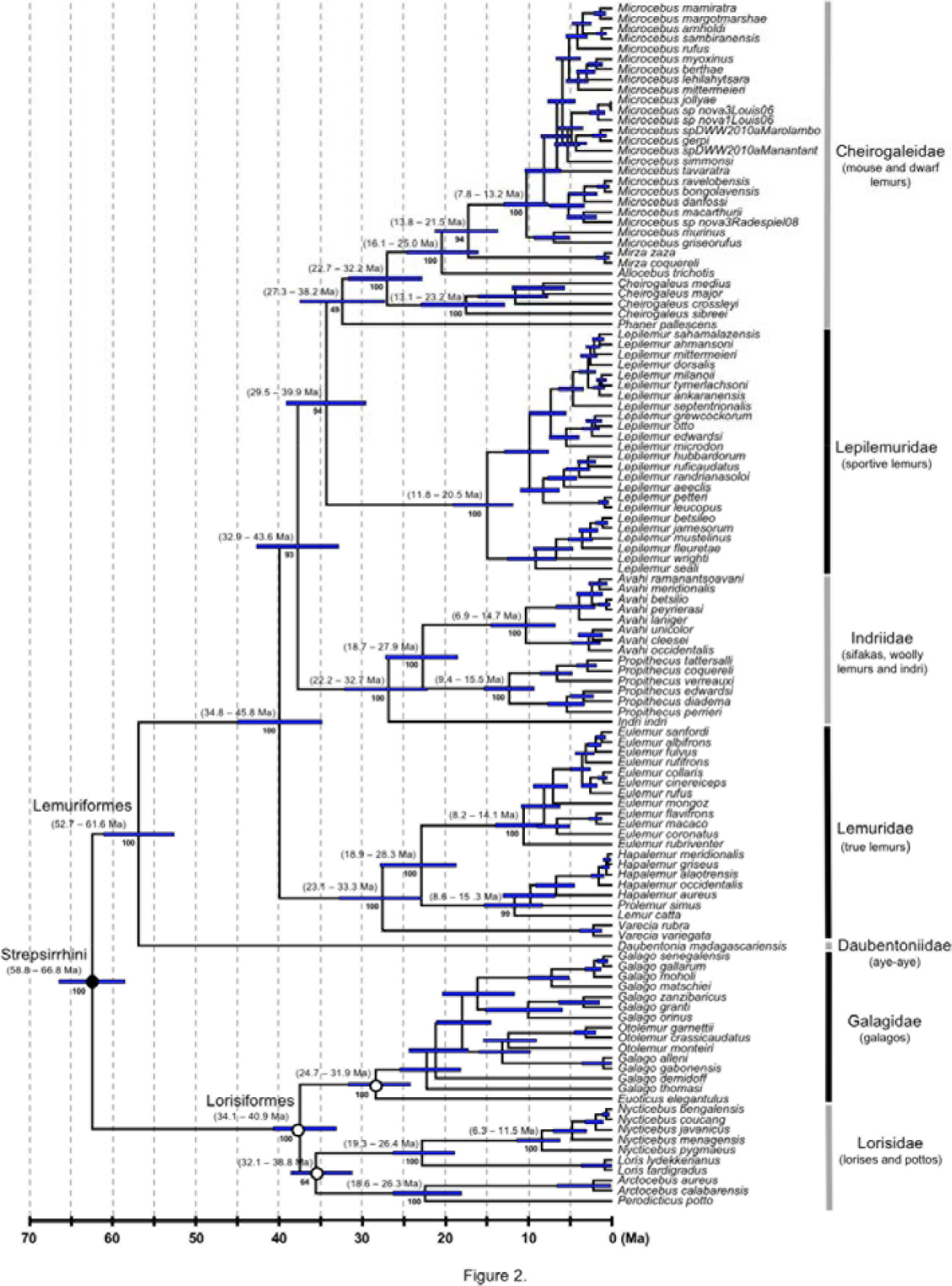
Strepsirrhine portion of the primate timetree (AR clock and calibration strategy A). Legend as for figure 1.

**Figure 3.**
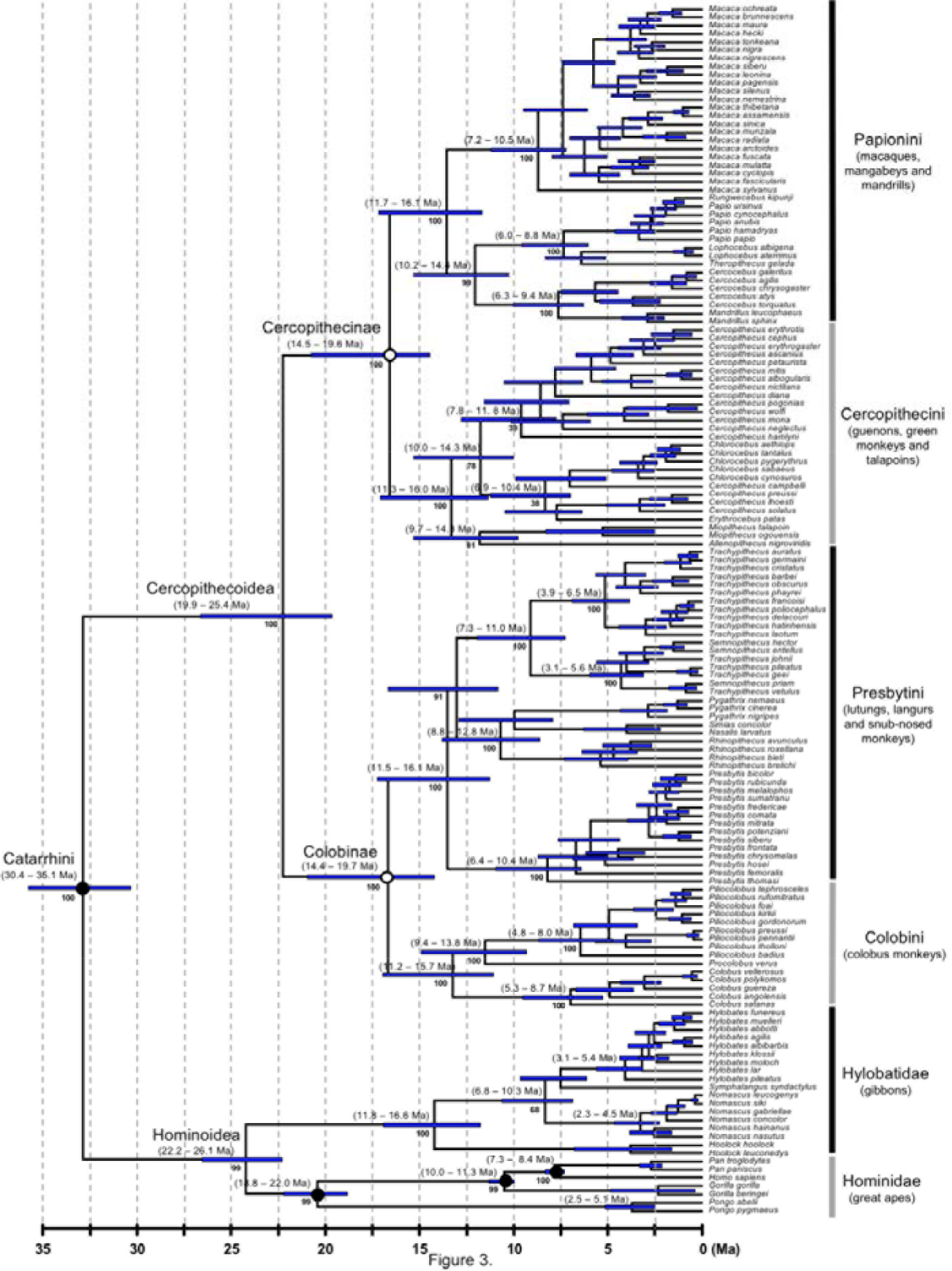
Catarrhine portion of the primate timetree [AR clock and calibration strategy A). Legend as for figure 1.

**Figure 4.**
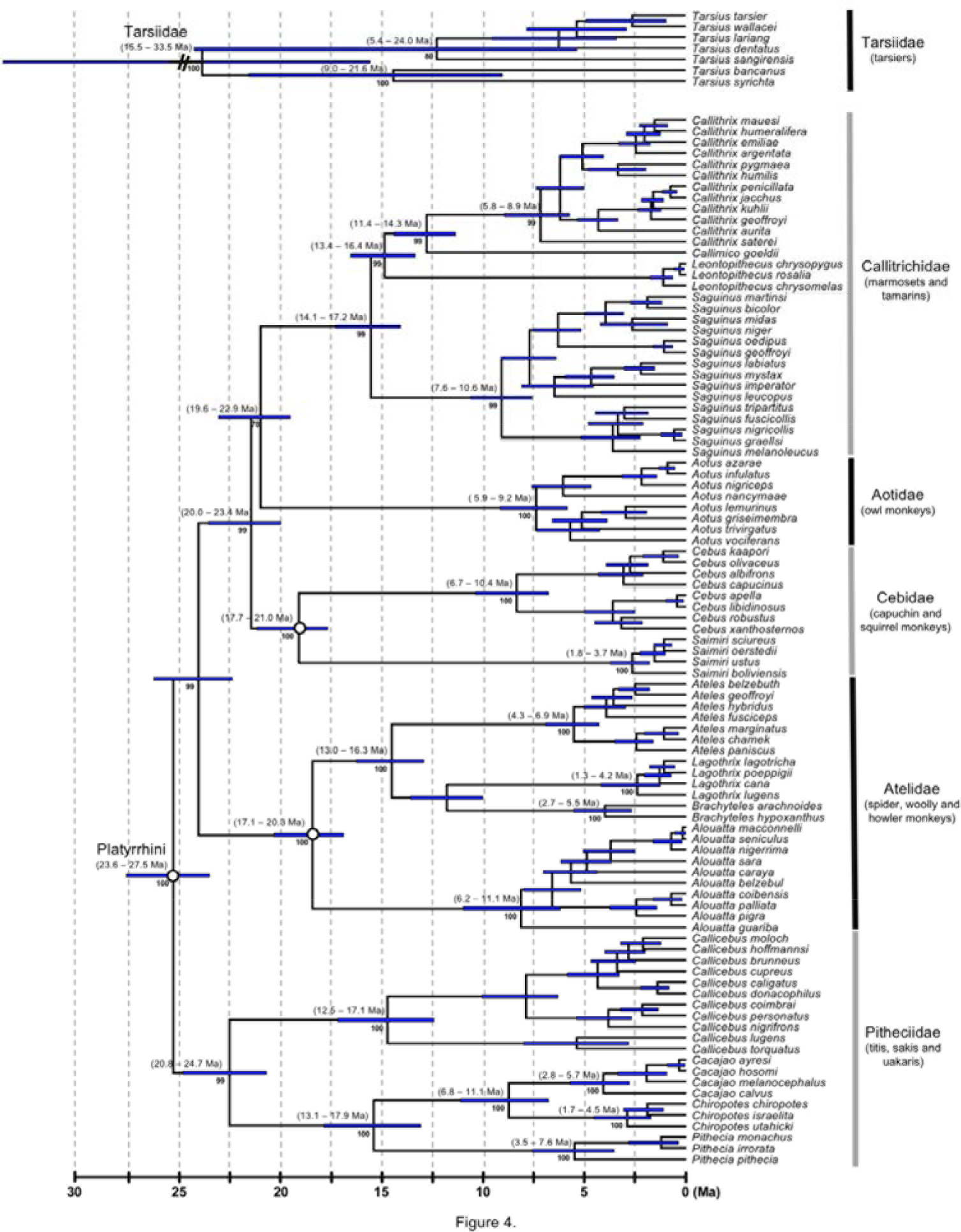
Tarsiidae and platyrrhine portion of the primate timetree [AR clock and calibration strategy A). Legend as for figure 1.

**Table 5.**
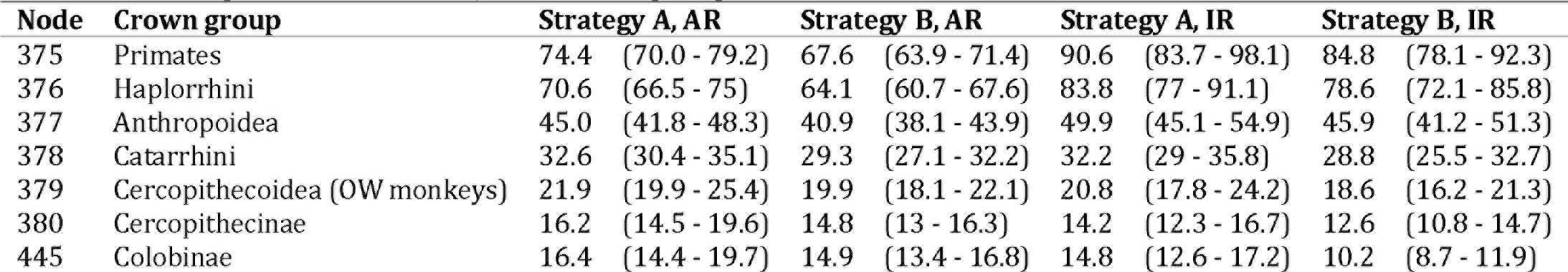
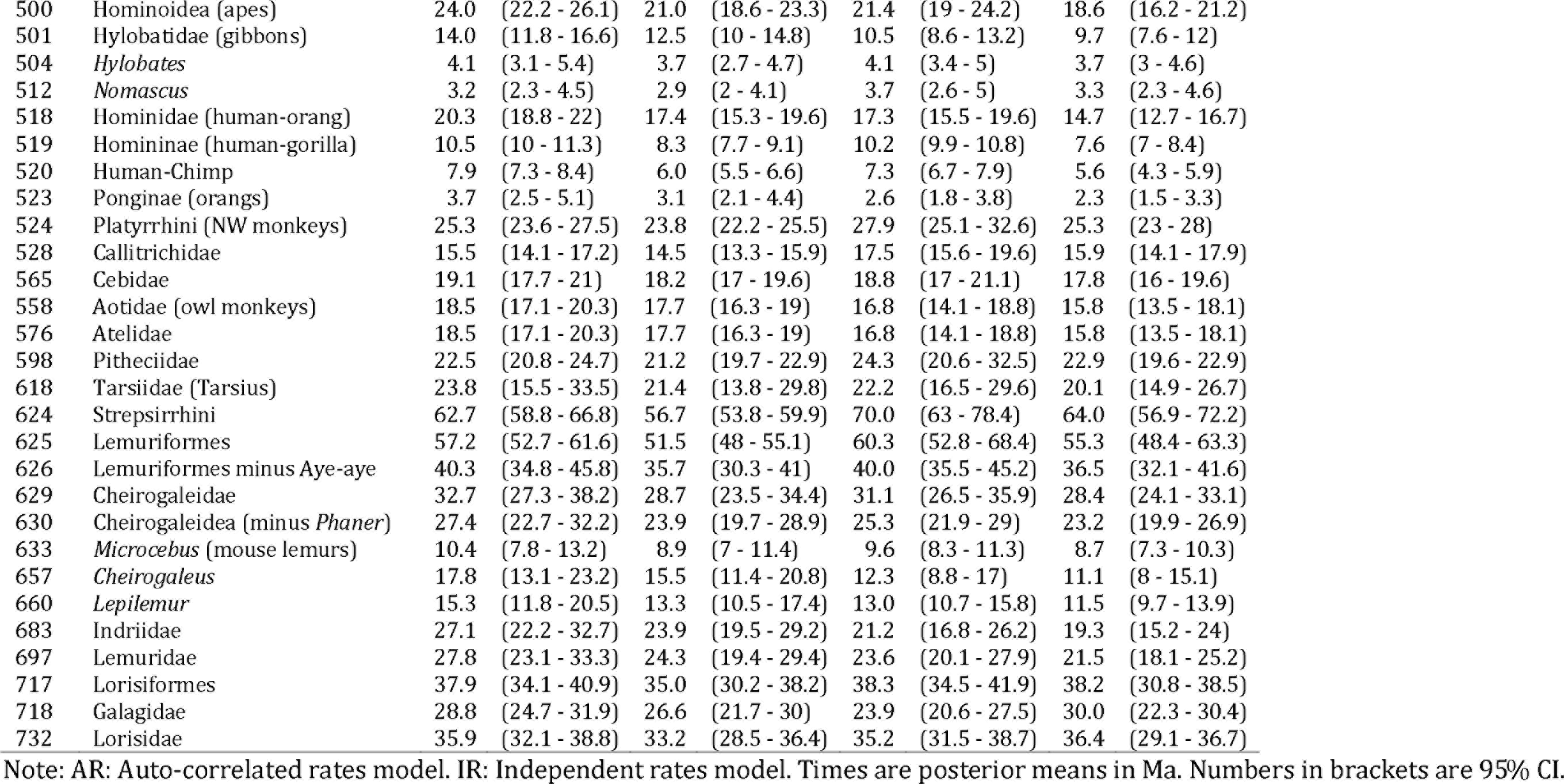
Divergence times of major Primate groups.

Calibration strategy B under the AR model gives similar node age estimates for the younger nodes in the tree (i.e. the 95% CI of node age overlap, Fig. 5 and Table 5). However, for the older nodes in the phylogeny (and in particular for Euarchonta, Primatomorpha, Primates, Haplorrhini, Lemuriformes and Lemuriformes minus aye-aye), strategy A produced older estimates (Figure 5). Under strategy B a pre-K-Pg origin of crown Primates is also favoured, although the posterior distribution of the age of crown Primates straddles the K-Pg boundary (71.4-63.9 Ma). The posterior probability for a pre-K-Pg origin of crown Primates is 80.0% under strategy B and 100% under strategy A.

**Figure 5.**
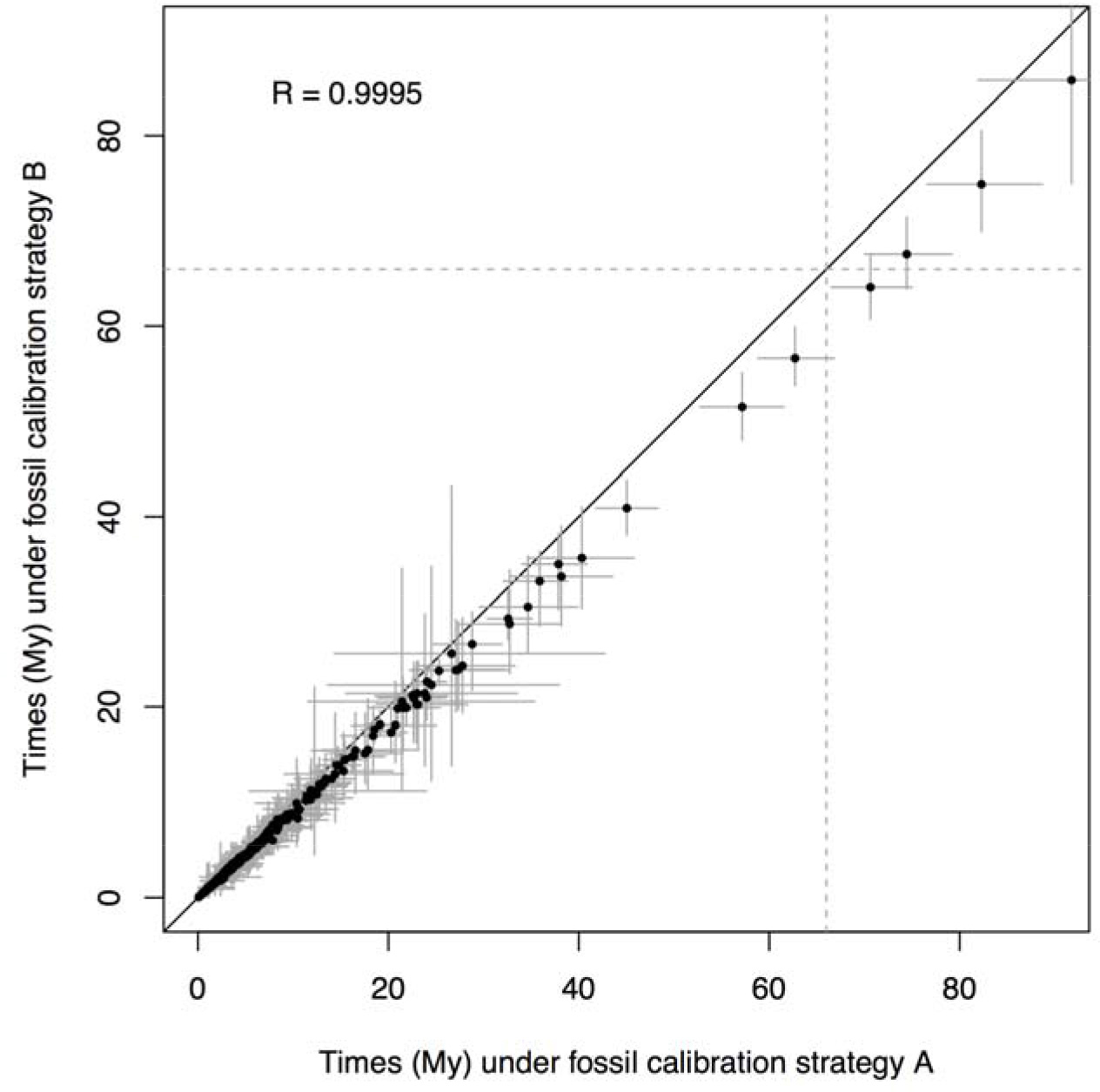
Posterior time estimates under fossil calibration strategy A vs. time estimates under strategy B, under the AR clock model.

Note that the two calibration strategies are in many cases based on the same fossils (Table 3), and the intervals defined by the fossil bounds overlap extensively between the two strategies. However, the seemingly small differences between the two strategies lead to noticeable differences in the posterior time estimates (Table 5, Fig. 5). In general, minimum bound constraints are older in strategy A than in strategy B (Table 3), and thus this may be the cause of the older time estimates in A vs. B. Constructing calibrations from uncertain fossil information is a subjective task, and different interpretations of the fossil record will lead to different posterior time estimates. The situation cannot be improved by adding more molecular data to the analysis, because the problem is statistically unconventional as we are trying to estimate two confounded parameters (times and rates) while the data are only informative about their product (the branch length). Furthermore, truncation effects among calibration densities affect construction of the time prior (Inoue, et al. 2010, Warnock, et al. 2015), and thus different calibration strategies may produce different time priors.

*Time prior and effect of truncation and outgroups*.- User-specified calibration densities usually do not satisfy the constraint that descendant nodes must be younger than their ancestors, thus the dating methodology must ‘truncate’ the calibration densities to satisfy the constraint to construct the time prior (Rannala, 2016). The result is that user-specified calibration densities and marginal priors of node ages may look substantially different. Figure 6 illustrates the effect of truncation on prior densities for strategies A and B. For example, in strategy B, the calibration densities on Euarchonta (the root of the phylogeny) and on Primates interact (the primate node has a Cauchy calibration density with a heavy tail), and consequently the prior density on the age of Euarchonta is pushed back (Fig. 6). The result is that the marginal prior age of Euarchonta ranges from 136-78 Ma (Fig. 6) instead of 130-61.5 Ma as in the calibration density (Table 3), while the upper age for the Primate prior is too old (127 Ma). In contrast, under strategy A, the calibration density on Primates has a much lighter tail, and thus the truncation effect with the Euarchonta node is minimal. The result is that the marginal time prior and the corresponding calibration densities for the Primates and Euarchonta nodes are very similar (Fig. 6). Similarly, under strategy B, the priors for two other nodes (Anthropoidea and Human-Gorilla) that use the heavy-tailed Cauchy calibrations have upper 95% limits that also appear unreasonably old (86.3 Ma and 25.0 Ma respectively). In general, calibration strategy A, which avoids using the long-tailed Cauchy calibrations, has calibration densities that are much closer to the resulting marginal priors, and thus strategy A results in a time prior which is much closer to the fossil information as interpreted by the palaeontologist.

**Figure 6.**
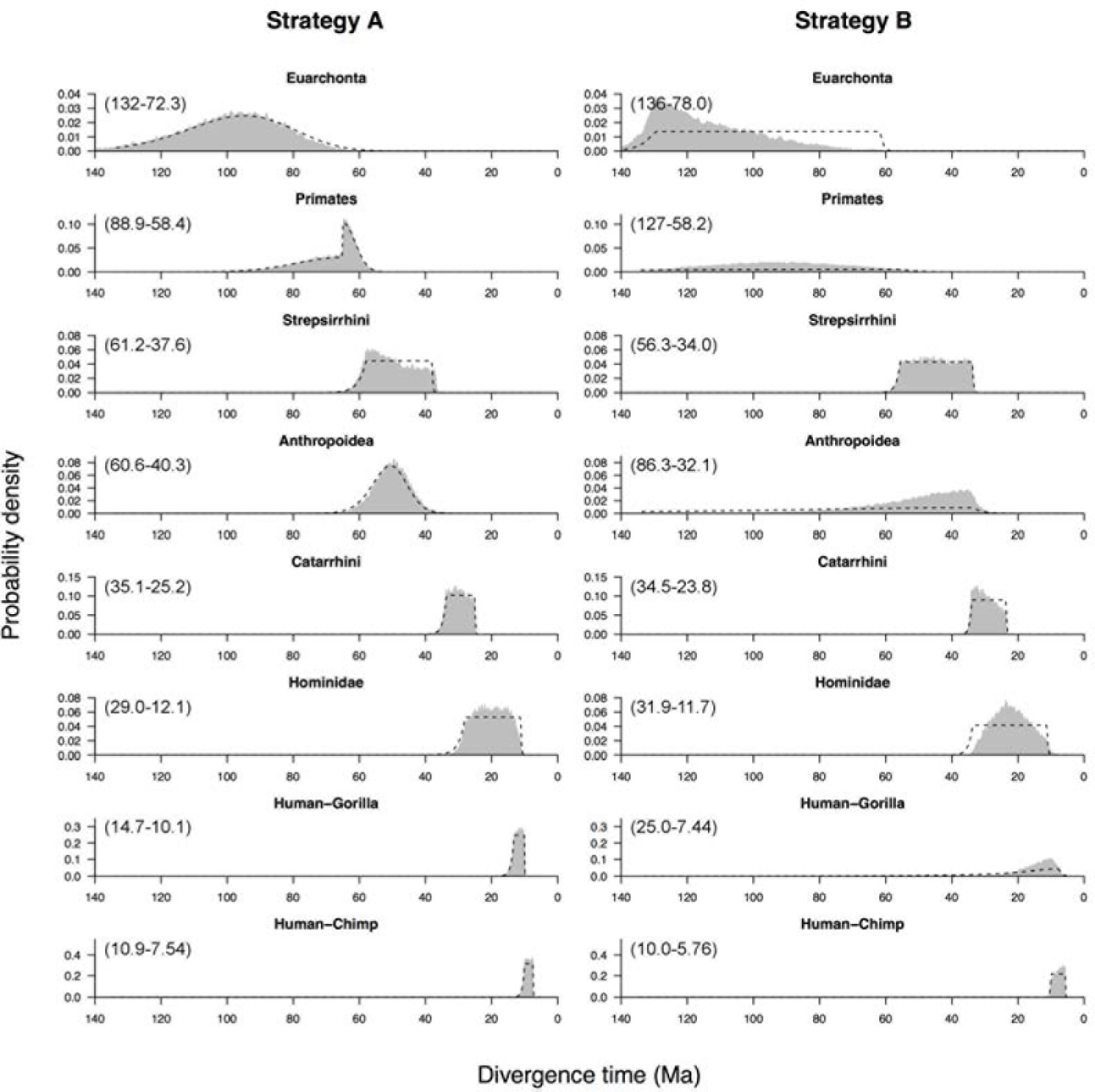
Calibration vs. prior densities for strategies A and B. Numbers in brackets indicate the 95% prior CL Note that the priors for three Cauchy-based calibrations in strategy B [Primates, Anthropoidea and Human-Gorilla) have heavy tails that extend substantially back in time.

*A Set of Calibrations for Mitogenomic Phylogenetic Analysis*–. Mitogenomic markers are widely used to construct phylogenies of closely related primate species with examples seen in phylogeographic studies of diversification of primates in the Amazon (Nascimento, et al. 2014) and in the timing of human diversification (Rieux, et al. 2014). The posterior distributions obtained here for the 10-species genomic data are useful calibrations for mitogenomic studies. Note that these cannot be used if the molecular alignment contains nuclear data as the calibrations already contain the information from nuclear genomes. The list of skew-t calibrations is provided in Table 6, together with approximations based on the gamma distribution, which can be used in software that does not implement skew-t calibrations (such as BEAST or MrBayes).

**Table 6.** Suggested skew-t and gamma calibrations for mitogenomic studies.

| **Crown group** | **Skew-t<sup>a,b</sup>** | **Gamma<sup>a,c</sup>** |
| --- | --- | --- |
| Primates | ST(0.878, 0.169, 2.41, 94.6) | G(78.6, 77.6) |
| Strepsirrhini | ST(0.580, 0.0567, 2.12, 149) | G(262, 327) |
| Haplorrhini | ST(0.705, 0.0628, 1.66, 960) | G(254, 411) |
| Anthropoidea | ST(0.415, 0.0291, 0.949, 294) | G(271, 363) |
| Catarrhini | ST(0.292, 0.0206, 0.995, 167) | G(316, 733) |
| Hominoidea | ST(0.185, 0.0167, 2.44, 312) | G(316, 1040) |
| Human-Gorilla | ST(0.0996, 0.0103, 19.6, 100) | G(311, 1575) |
| Human-Chimp | ST(0.0788, 0.00687, 3.51, 6.15) | G(292, 2715) |

a. Densities calculated under fossil calibration strategy A and AR model using a time unit of 100 My.
b. The parameters of the Skew-t distribution are location, scale, shape and df.
c. Note that here we use the shape ( $\alpha$ ) and rate ( $\beta$ ) parameterization. For the scale parameterization use $s = 1/\beta$ . The mean is $\alpha/\beta$ and variance is $\alpha/\beta^2$ .

### Effect of the Clock Model

In the auto-correlated clock (AR) model, rates for branches are assumed to evolve according to a geometric Brownian diffusion process, where the rate for a particular branch depends on the rate of its parent branch. Large rate changes from an ancestral branch to its daughter branches are penalized. In the independent rate (IR) model, rates for branches are assumed to evolve independently among branches, and sharp rate changes from parent to daughter branch are not penalized. Posterior means and 95%
CIs for locus (partition) rates obtained under both clock models are given in Table 7. The clock model has a strong impact on posterior time estimates, particularly for the most ancient nodes in the phylogeny. Under the IR model, the ages of Euarchonta, Primatomorpha, Primates, Haplorrhini and Strepsirrhini are substantially older than those estimated under the AR model (Table 5 and Figure 7).

**Table 7.** Posterior means of locus rates and rate variance parameters under calibration strategy A.

| **Partition** | **AR locus mean<sup>a</sup>** | **AR locus $\sigma^2$** | **IR locus mean<sup>a</sup>** | **IR locus $\sigma^2$** |
| --- | --- | --- | --- | --- |
| 1. mit 1st+2nd | 0.20 (0.086–0.38) | 1.6 (1.1–2.2) | 0.37 (0.33–0.41) | 0.33 (0.26–0.41) |
| 2. mit 3rd | 1.8 (0.63–3.7) | 4.6 (3.6–5.8) | 3.0 (2.7–3.3) | 0.24 (0.19–0.30) |
| 3. mit RNA | 0.53 (0.23–1.0) | 1.8 (1.1–2.7) | 0.54 (0.48–0.60) | 0.31 (0.23–0.41) |
| 4. nuclear 1st+2nd | 0.11 (0.041–0.24) | 2.4 (1.6–3.3) | 0.037 (0.033–0.041) | 0.37 (0.27–0.51) |
| 5. nuclear 3rd | 0.34 (0.13–0.71) | 2.3 (1.6–3.1) | 0.11 (0.098–0.12) | 0.33 (0.23–0.47) |
| 6. nuclear non-coding | 0.20 (0.065–0.47) | 3.3 (2.4–4.4) | 0.068 (0.061–0.075) | 0.40 (0.30–0.56) |
| 7. nuclear 1st+2nd | 0.038 (0.024–0.057) | 0.26 (0.12–0.51) | 0.027 (0.022–0.034) | 0.14 (0.062–0.28) |
| 8. nuclear 3rd | 0.16 (0.099–0.23) | 0.27 (0.12–0.54) | 0.11 (0.090–0.14) | 0.17 (0.077–0.33) |

a. Substitutions per site per 100 My. For example 0.21 means $2.1 \times 10^{-9}$ subs/site/year.

**Figure 7.**
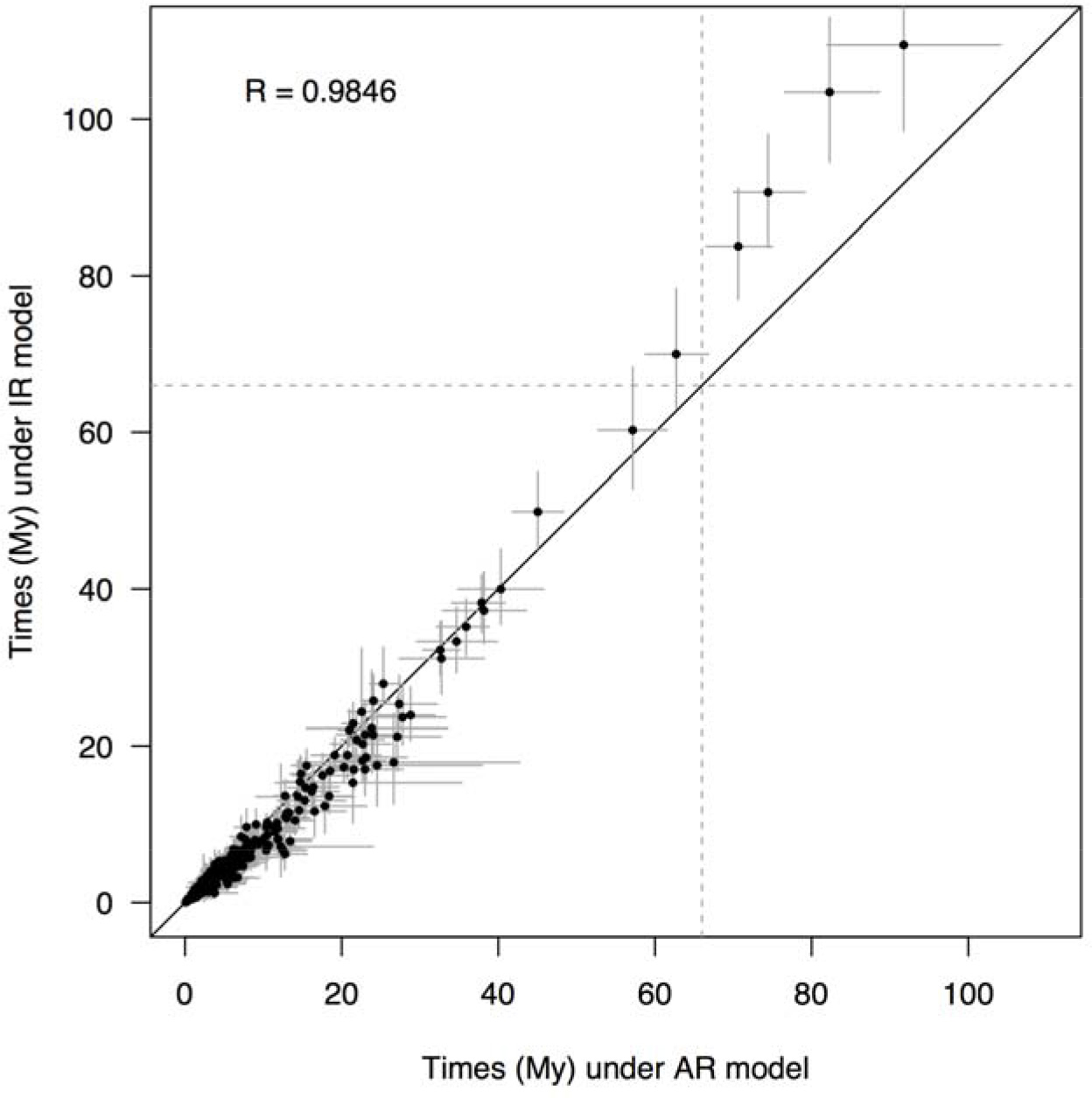
Posterior time estimates under the AR vs. IR clock models, for calibration strategy A.

Results of Bayesian model selection of clock model using thermodynamic integration are shown in Table 8. The AR model has the highest marginal likelihood in 5 out of the 6 partitions analysed, with the posterior model probability > 90% in two partitions, and 79%, 66%, 53% and 29% in the other four. When the six partitions are analysed in a multi-partition dataset, the posterior probability is virtually 100% in favour of the AR model. We note that ideally, the marginal likelihood calculations should have been carried out on the complete dataset, but unfortunately, this is so computationally expensive that it cannot be done in a feasible amount of time. Although the estimates are based on the data subset, it appears unlikely that the results would change for the whole data, given the consistent support for the AR model.

**Table 8.** Bayesian model selection of rate model.

| **Dataset** | **Model** | **Log Marginal $L^a$** | **BF** | **$P^b$** |
| --- | --- | --- | --- | --- |
| Mitochondrial 1<sup>st</sup> and 2<sup>nd</sup> c.p. | SC | -16,519.03 (0.010) | $1.3 \times 10^{-18}$ | $1.2 \times 10^{-18}$ |
|  | IR | -16,480.58 (0.021) | 0.063 | 0.060 |
|  | **AR** | **-16,477.82 (0.035)** | - | **0.94** |
| Mitochondrial 3<sup>rd</sup> c.p. | **SC** | **-16,684.50 (0.014)** | - | **0.61** |
|  | IR | -16,686.29 (0.043) | 0.17 | 0.10 |
|  | AR | -16,685.26 (0.040) | 0.47 | 0.29 |
| Mitochondrial RNA | SC | -7,906.85 (0.0087) | 0.74 | 0.39 |
|  | IR | -7,908.40 (0.015) | 0.16 | 0.08 |
|  | **AR** | **-7906.55 (0.023)** | - | **0.53** |
| Nuclear 1<sup>st</sup> and 2<sup>nd</sup> c.p. | SC | -32,179.80 (0.0092) | 0.0047 | 0.0037 |
|  | IR | -32,175.77 (0.022) | 0.27 | 0.21 |
|  | **AR** | **-32,174.44 (0.032)** | - | **0.79** |
| Nuclear 3<sup>rd</sup> c.p. | SC | -24,535.33 (0.012) | $7.2 \times 10^{-12}$ | $6.7 \times 10^{-12}$ |
|  | IR | -24,512.45 (0.038) | 0.062 | 0.058 |
|  | **AR** | **-24,509.67 (0.030)** | - | **0.94** |
| Nuclear UTR and introns | SC | -64,739.20 (0.016) | $5.7 \times 10^{-4}$ | $3.8 \times 10^{-4}$ |
|  | IR | -64,732.41 (0.038) | 0.51 | 0.34 |
|  | **AR** | **-64,731.73 (0.046)** | - | **0.66** |
| All<sup>c</sup> | SC | -162,684.8 (0.024) | $2.1 \times 10^{-103}$ | $2.1 \times 10^{-103}$ |
| | IR | -162,467.0 (0.086) | $8.4 \times 10^{-9}$ | $8.4 \times 10^{-9}$ |
|  | **AR** | **-162,448.4 (0.15)** | - | **1.00** |

Marginal likelihoods are estimated by thermodynamic integration with 64 points. The substitution model is HKY+G. SC: Strict clock; IR: Independent log-normal; AR: Auto-correlated rates. The age of the root is fixed to one (i.e. we use a 'B(0.99, 1.01)' calibration on the root in MCMCTree). The rate priors are G(2, 1) and G(2, 20) for mitochondrial and nuclear data respectively. The prior on $\sigma^2$ is G(1, 1) in all cases. The model with the highest posterior probability in each dataset is shown in bold type. a: Values in brackets are the standard errors (see Appendix 2). b: Posterior model probabilities are calculated assuming a uniform prior on models. c: The six datasets are analysed together as six partitions.

### Effect of genome-scale data

Figure 8 shows the infinite-sites plot for the primate data analysed here. For calibration strategy A, the eight primate nodes shared between the 10-species and 372-species trees (i.e. the nodes constrained by the large genome-scale alignment, table 2) fall in an almost perfectly straight line (*R* = 0.992, Fig. 8A). This indicates that for these nodes, uncertainty in the time estimates is dominated by uncertainties in the fossil calibrations rather than by uncertainties in the molecular data. For strategy A, a regression line through the origin fitted to the eight data points is *w* = 0.128*t*, meaning that for every 1 My of divergence, 0.128 My are added to the CI width (Fig. 8A). On the other hand, when considering all 371 nodes in the tree, the relationship between CI width and mean times is far from linear, and the level of uncertainty is much higher. In this case, 0.277 My are added to the CI width for every 1 My of divergence. The trend is similar under calibration strategy B (Fig. 8B), albeit in this case there is in general more uncertainty in time estimates (i.e. the slope of the regression lines is larger). This appears due to strategy B being more conservative than strategy A, that is, some of the calibration densities used in B are substantially wider, encompassing larger time ranges (Fig. 6).

**Figure 8.**
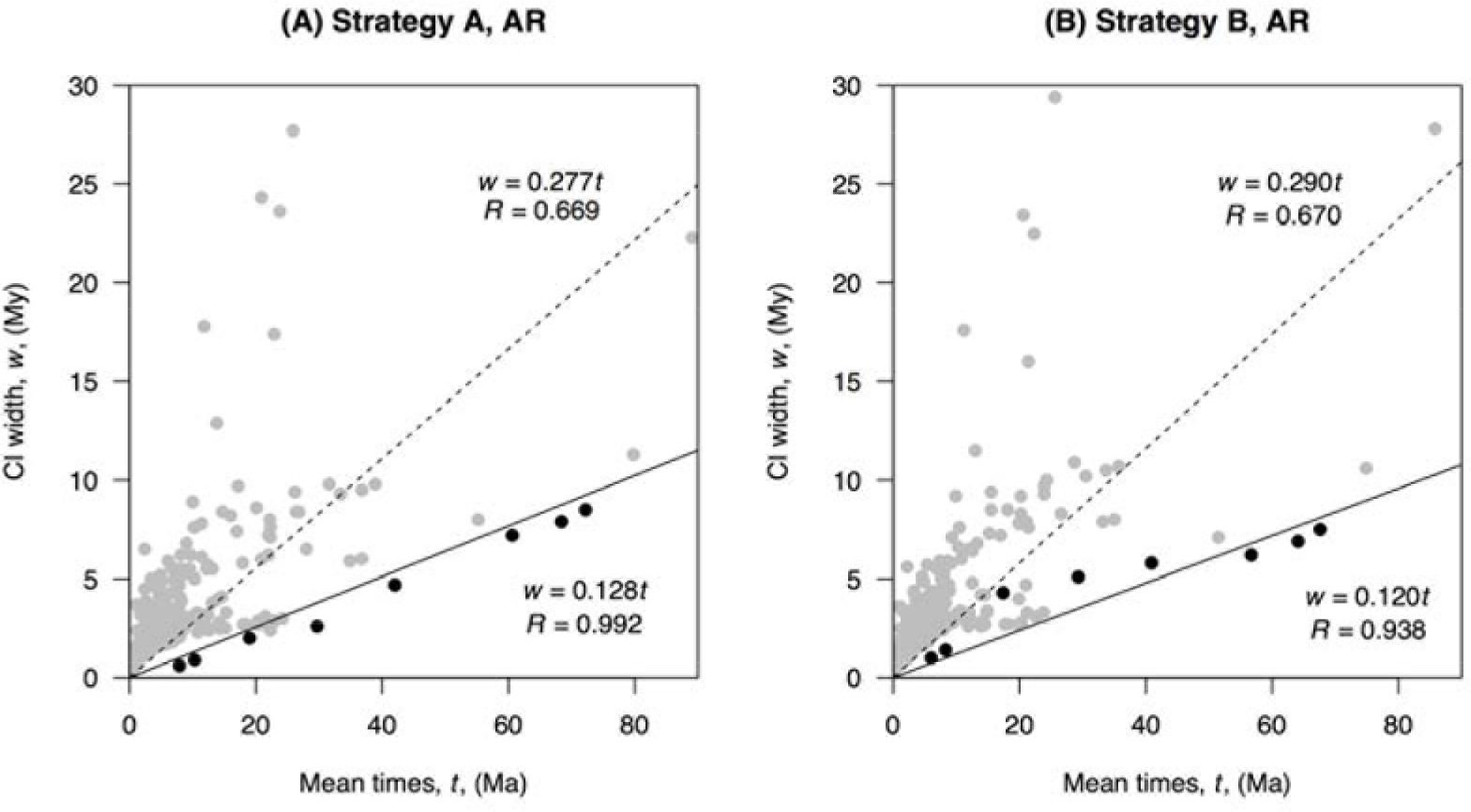
Infinite-sites plot Posterior CI width is plotted against mean posterior divergence times for (A) analysis under calibration strategy A and AR clock, and (B) analysis under calibration strategy B and AR clock. In both cases, black dots indicate the eight primate nodes shared between the 10-species and 372-species trees, while the grey dots represent the rest of the nodes. Solid line: regression through the origin fitted to the black dots. Dashed line: regression through the origin fitted to all the dots.

## DISCUSSION

### A Phylogenomic View of Primate Divergences

A primary aim of this work was to study the effect of genome-scale data on divergence time estimates on a species-level phylogeny. Given the wide availability of whole genome data for a core-set of species, it is important to know whether the use of these data for a subsample of lineages will be enough to reduce time uncertainties in a species-level phylogeny to the theoretical limit. The results of figure 8 clearly indicate this is not the case. Although for the core ancestral nodes in the Primate phylogeny, the genome-scale alignments do constrain uncertainty in time estimates close to their theoretical limit (so it is highly unlikely that adding additional molecular data for these species will improve time estimates appreciably), for species without genome-scale data, there are still substantial uncertainties left for family-level and genus-level divergences in the tree. For some nodes, the CI-width is almost as large as the node age (for example, for Tarsiidae, the node age is 23.8 Ma with CI-with 18 My, which is 76% of the node age, table 5). Thus much work is still needed in order to improve time estimates for the hundreds of more recent divergences in the tree. Furthermore, application of morphological-based models for dating (Ronquist et al. 2012) and the fossilised birth-death process (Heath et al. 2014) also offer exciting prospects and challenges in obtaining time estimates for the species-level divergences (O’Reilly et al. 2015, dos Reis et al. 2016). Improving these estimates will be important in studies of primate diversification rates and to correlate primate diversification events with major geological events in the history of the Planet (such as glaciations, continental drift, the closure the Panama isthmus, etc.).

### Sequential Bayesian Analysis verus Secondary Calibrations

In this work we used the posterior of times obtained under a small dataset as the prior of times in a second analysis under a large dataset. This approach is justified as long as the datasets are independent under the likelihood model and as long as the datasets do not overlap (that is, they share no genes). The use of the posterior in an analysis as the prior for the next is a well-known feature of Bayesian inference (Gelman et al. 2013). Consider data that can be split into two subsets, *D* = (*D*_1_, *D*_2_), which are independent under the likelihood model. The posterior distribution for parameter *θ* is

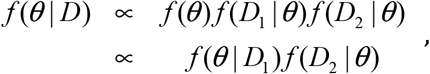

where *f*(*θ* | *D*_1_) ∝ *f*(*θ*) *f*(*D*_1_ | *θ*) is the posterior distribution of *θ* when only *D*_1_ are analysed. It is apparent that using *f*(*θ* | *D*_1_) as the prior when analysing *D*_2_ leads to the posterior for the joint data *D*. In other words, performing the analysis in one step (joint analysis of *D*_1_ and *D*_2_) or in two steps (posterior under *D*_1_ as prior under *D*_2_) results in the same posterior distribution.

The approach we used here to analyse the primate data is justified because the likelihood model assumes that the sequence partitions are non-overlapping and independent. However our approach is approximate. In multi-parameter models, the posterior is a multidimensional distribution that may have a complex correlation structure. Here we ignored the correlation structure of the nine times estimated using the genomic data, and approximated the corresponding high-dimensional time posterior as the product of the marginal densities of the times, although a truncation is applied to ensure that descendants are younger than ancestors. Note that joint analysis of all the partitions would have been preferable, but it is computationally prohibitive.

This Bayesian sequential analysis is different from the use of secondary calibrations in some dating studies (Graur and Martin, 2004), where the secondary calibrations were used as point calibrations (with the uncertainties on node age estimates ignored), and where in many cases the data analysed under the secondary calibration was the same as the data analysed to obtain the calibration in the first place. We stress that the Bayesian sequential approach is justified only if the data subsets do not overlap. The genes/sequences analysed in the first step must be different from those analysed in the second step.

### Clock Model

An interesting result from our study is the finding that the AR model fits the primate data better than the IR relaxed-clock model. In the context of previous studies, Lepage, et al. (2007) found, using Bayes factors and no fossil calibrations, that two AR relaxed clocks (CIR and log-normal) fitted real data (eukaryotes, mammals and vertebrates) better than IR models. More recently, Lartillot, et al. (2016) introduced a mixed relaxed clock that has auto-correlated- and independent-rates components. In their analysis, the mixed clock appeared to provide a better description of rate evolution in the mammal phylogeny, however, they did not assess clock model fit with Bayes Factors. Linder et al. (2011) found, also by using Bayes factors, that IR models better fit an angiosperm phylogeny better than AR models. Additionally, they found that, when analysed without fossil calibrations, the AR model fit an ape phylogeny better than the IR model. However, when analysed with fossil calibrations, the IR model fit the ape data better.

In the AR model the variance of the log-rate for branches is proportional to the time of divergence, so that the variance is expected to be close to zero for closely related species. In other words, the AR model allows for “local clocks” for closely related species, while allowing the rate to drift substantially across distantly related clades. This model is, from a biological point of view, quite appealing intuitively, and it also fits anecdotal evidence where the strict clock cannot be statistically rejected among very closely related species, for example, among the apes (dos Reis, et al. 2016, box 2). In contrast, the IR model assumes that the variance of the branch rates is the same for different time scales. This would appear biologically unrealistic. Arguments have been put forward in favour and against both of the two types of relaxed-clock models examined by our study (Thorne et al. 1998, Drummond et al. 2006, Ho 2009), and clearly further research is still needed to understand which clock model is the most biologically realistic and appropriate for real data analysis. This will be a challenging task given how difficult it has been to distinguish between the two models in simulation studies (Heat et al. 2012, Ho et al. 2015, Lepage et al. 2007).

### Five Decades of Primate Molecular Timetrees

The project of employing genomic information for discovering the geological age of the Primates began virtually simultaneously with the publication of Zuckerkandl and Pauling's (1965) molecular clock hypothesis. Sarich and Wilson (1967) employed a strict clock interpretation of immunological distance data to hypothesize that humans and other African apes (i.e., chimp and gorilla) shared a common ancestor as recently as five million years ago. This was revolutionary at the time given the implications for the necessarily rapid evolution of bipedal locomotion in the hominin lineage, and accordingly, drew considerable attention from anthropological community (Read and Lestrel, 1970; Uzzell and Pilbeam, 1971; Lovejoy et al., 1972; Radinsky, 1978; Corruccini et al, 1980). Despite this interest, it wasn't until the 1980s that the field of divergence time estimation assumed a relatively modern flavour. It was only then that investigators began to apply statistical models to DNA sequence data for the purposes of branch length and divergence time estimation (e.g., Hasegawa et al., 1985). Remarkably, these studies first emerged at a time when the sister-lineage relationship of humans to chimps was considered highly controversial (e.g., Goodman et al., 1983) — a relationship that is now considered unequivocal.

From this point forward, primate timetrees have been produced with increasing frequency, though with widely varying conclusions regarding the age of the last common ancestor of Primates and of the major subclades within the Order (Table 1). Prior to the study reported here, the estimated age of the living primate clade has spanned a 30 My differential, ranging from as young as 55 Ma (Koop et al., 1989) to as old as 87 Ma (Perelman et al, 2011). The new millennium has been a particularly active time for primate divergence time analysis. Beginning in the early 2000's, published studies have employed a diverse assortment of datasets applied to the problem (e.g., nuclear, mitochondrial, and their combination), as well as a range of statistical methods and calibration densities. Despite this array of data and methods, all of these studies—with only one notable outlier (Chatterjee, 2009)—have consistently indicated that the crown Primates clade originated prior to the K-Pg event (see also Steiper and Seiffert, 2012). Given the continued dearth of fossil data to support this hypothesis, however, the result continues to be viewed with scepticism by the paleoanthropological community (Bloch et al., 2007; Silcox, 2008; O’Leary et al., 2013; but see Martin et al., 2007).

As described at length above, the current study gives added weight to the conclusion that primates are an ancient clade of placental mammals, arising just prior to or millions of years before the K-Pg. And even though lineage diversification within the major subclades is hypothesized not to have occurred until after the commencement of the Paleogene, the separation of tarsiers from other haplorrhines, and the divergence of haplorrhines and strepsirrhines, consistently appear to proceed or nearly coincide with the K-Pg. Given that this event was unequivocally one of the most disruptive and destructive geological episodes in Earth history, the temporal coincidence speaks both to the ecological flexibility and to the evolutionary opportunism of the earliest primates. Although now extinct in North America and in Europe, the primate fossil record shows that the clade was once nearly pan-global, even potentially including Antarctica. Thus, when viewed in the context of divergence date estimates, all of which fall within a temporal window when, as now, continental and island residences would have already been sundered by significant oceanic barriers (most notably, the separation of South America from Africa by the Atlantic Ocean), we must conclude that early primates would have been able dispersers. In fact, the ability to cross barriers, both terrestrial and aquatic, and to successfully colonize new land masses, are distinct hallmarks of the primate radiation (Gingerich, 1981; Yoder and Nowak, 2006; de Queiroz, 2005, 2014; Seiffert, 2012; Beard, 2016; Bloch et al., 2016).

## SUPPLEMENTARY MATERIAL

Data available from the Dryad Digital Repository: http://dx.doi.org/10.5061/dryad. [NNNN]

## DEDICATION

This paper is dedicated to the memory of our co-author Gregg F Gunnell.

## APPENDIX 1

Justifications for dates assigned to 17 fossil calibrations in this study (Table 3) are given below: 8 calibrations for strategy A (SA), 1 calibration for strategy B (SB), and 8 calibrations shared among both strategies (SAB). The justifications for the remaining calibrations in Table 3 are given in dos Reis et al. (2012; see also Benton et al. 2009). In some cases, the dates used are not exactly those published in cited references. In these cases, the dates utilized reflect published as well as unpublished information or adjustments deemed necessary given the uncertainty of some dates. In any case the discrepancies are always small and unimportant considering the breadth of the fossil calibrations. Note that specifying maximum bounds is a difficult task, in particular because absence of fossil evidence is not evidence that a clade did not exist in a point in time (Ho and Philips, 2009). We use stem fossils as benchmark points onto which to construct diffuse maximum bounds (i.e. with a large probability of violation, *p*_*U*_) on some node ages.

## Hominini | *Homo-Pan* | 7.5 Ma – 10 Ma | SA

The minimum age for the divergence of hominins is placed at 7.5 Ma and is based on the appearance of †*Sahelanthropus* at 7.2 Ma (Brunet, et al. 2002, Brunet, et al. 2005, Lebatard, et al. 2008). There is some controversy as to the proper taxonomic position of *Sahelanthropus* (Wolpoff, et al. 2006, MacLatchy, et al. 2010) but we regard it as the oldest record of a plausible crown hominin. *Sahelanthropus* comes from the Anthracotheriid Unit of an unnamed formation in the Mega-Chad Basin in Chad (Brunet, et al. 2005). The associated mammalian fauna is very similar to that found from the Nawata Formation, Lothagam, Kenya which maybe as old as 7.4 Ma (MacDougall 2003). The divergence of the hominin lineage seems unlikely to have occurred before the appearance of the potential gorillin *Chororapithecus* at 10 Ma (Suwa, et al. 2007, Harrison 2010a).

## Homininae | *Gorilla-Homo* | 10 Ma – 13.2 Ma | SA

The minimum age for the divergence of crown hominines is placed at 10 Ma based on the appearance of the potential gorillin *Chororapithecus*. Like *Sahelanthropus*, the taxonomic status of *Chororapithecus* is not without controversy (Suwa, et al. 2007, Harrison 2010a). Harrison (2010a) regards *Chororapithecus* as best interpreted as a stem hominin or even a stem hominid – we feel that the features that do support a relationship with gorillas are well enough established to use the date of appearance of *Chororapithecus* as a minimum divergence date for hominines. *Chororapithecus* comes from the late Miocene Beticha section of the Chorora Formation in Ethiopia and is dated at 10-10.5 Ma (Geraads, et al. 2002). We use a maximum divergence date of 13.2 (Raza, et al. 1983) for *Sivapithecus* but it is now evident that this date might be slightly too old. In light of this the divergence of the hominine lineage is unlikely to have taken place before the earliest appearance of the probable crown pongine *Sivapithecus* at 12.5 Ma (Begun 2010, Begun, et al. 2012).

## Hominidae | *Pongo-Homo* | 11.2 Ma – 28 Ma | SA

The minimum age for the divergence of crown hominids is placed at 11.2 Ma based on the earliest appearance of the crown pongine *Sivapithecus* (Kappelman, et al. 1991, Begun 2010, Begun, et al. 2012). *Sivapithecus* is known from Siwalik group rocks (Chinji, Nagri and Dhok Pathan formations) in Indo-Pakistan that range in age from 14 Ma to 5.5 Ma (Badgley and Behrensmeyer 1995) with *Sivapithecus* restricted to a range of 12.5 Ma to 7.4 Ma (Flynn, et al. 1995). The divergence of crown hominids is unlikely to have occurred before the first appearance of *Kamoyapithecus* at 25 Ma (Seiffert 2010). We use 28 Ma as a slightly more conservative maximum.

## Catarrhini | *Homo-Macaca* | 25 Ma – 33.7 Ma | SA

The presence of the crown hominoid *Kamoyapithecus* (Zalmout, et al. 2010) indicates that the minimum divergence time for crown Catarrhini is 25 Ma (Seiffert 2010). *Kamoyapithecus* is only known from the Erageliet beds, Kalakol Basalts locality of Lothidok in Kenya (Madden 1980, Leakey, et al. 1995, Rasmussen and Gutierrez 2009). A soft maximum of 33.7 Ma on the age of crown catarrhines is given due to the absence of hominoids before 33.7 Ma.

## Anthropoidea | Catarrhini-Platyrrhini | 41 Ma – 62.1 Ma | SA

Calibration density constructed from fossil modeling. The effects of the K-Pg extinction are included in the model. See Wilkinson, et al. (2011) for details.

## Haplorrhini | crown *Tarsius* | 45 Ma | SB

The presence of the crown tarsiid *Xanthorhysis* in Shanxi Province, China (Beard 1998) and apparently of the genus *Tarsius* in fissure fills at Shanghuang in Jiangsu Province, China (Beard, et al. 1994), both dating to the late middle Eocene (40-45 Ma), circumscribe the minimum divergence time for crown Haplorrhini at 45 Ma.

## Strepsirrhini | Lorisiformes-Lemuriformes | 37 Ma – 58 Ma | SA

The minimum age for the divergence of crown Strepsirrhini is placed at 37 Ma based on the first appearance of the crown lorisiform *Saharagalago* (Seiffert, et al. 2003). *Saharagalago* is only known from Fayum Quarry BQ-2 in the Birket Qarun Formation, Egypt. The divergence of crown strepsirrhines is unlikely to have occurred before the first appearance of the basal primate *Altiatlasius* from Ouarzazate in Morocco which is considered to represent the late Paleocene (Thanetian) and dating to around 58 Ma (Gheerbrant, et al. 1993, Gheerbrant, et al. 1998, Seiffert 2010).

## Primates | Haplorrhini-Strepsirrhini | 57.6 Ma – 88.6 Ma | SA

Calibration density constructed from fossil modeling. Includes the effects of the K-Pg extinction in the model. See Wilkinson, et al. (2011) for details.

## Euarchonta | Scandentia-Primates | 65 Ma – 130 Ma | SA

The minimum age for the divergence of crown Euarchonta is placed at 65 Ma based on the first appearance of the crown euarchontan *Purgatorius* (Bloch, et al. 2007). *Purgatorius* is known from the early Paleocene (Puercan) Tullock and Bear Formations in Montana (Clemens 1974, Buckley 1997, Clemens 2004, Chester, et al. 2015) and from the earliest Paleocene Ravenscrag Formation in Saskatchewan (Fox and Scott 2011). The divergence of Euarchonta is unlikely to have been before the appearance of placental mammals by at least 130 Ma (Luo 2007, but see Luo, et al. 2011 for a potential 130 My old eutherian).

## Lorisidae | *Nycticebus-Perodicticus* | 14 Ma – 37 Ma | SAB

The minimum age for the divergence of crown Lorisidae is placed at 14 Ma based on an undescribed genus and species from Fort Ternan in Kenya cited by Walker (1978) and Harrison (2010b). The minimum age could possibly be as old as 19 Ma if *Mioeuoticus* (Leakey in Bishop 1962, Walker 1978) represents a crown lorisid (Harrison 2010b). Fossil lorisids are known from the early to middle Miocene in Africa (Phillips and Walker 2000, 2002, Harrison 2010b) and from the late Miocene of Pakistan (Jacobs 1981). The divergence of lorisoids is unlikely to have occurred before the first appearance of the potential stem lorisid *Karanisia* at BQ-2 in Egypt (Seiffert, et al. 2003).

## Galagidae | *Galago-Euoticus* 115 Ma – 37 Ma | SAB

The minimum age for the divergence of crown Galagidae is placed at 15 Ma based on an undescribed genus and species from Maboko Island in Kenya cited by McCrossin (1999) and Harrison (2010b). The minimum age could possibly be as old as 19 Ma if either *Progalago* or *Komba* represent a crown galagid (MacInnes 1943, Simpson 1967, Harrison 2010b). Fossil galagids are known from the early Miocene through early Pleistocene in Africa (Phillips and Walker 2002, Harrison 2010b). The divergence of galagids is unlikely to have occurred before the first appearance of the potential stem lorisid *Karanisia* at BQ-2 in Egypt (Seiffert, et al. 2003).

## Lorisiformes | *Galago-Perodicticus* | 18 Ma – 38 Ma | SAB

The minimum age for the divergence of crown Lorisiformes is placed at 18 Ma based on the appearance of the potential crown lorisoid *Mioeuoticus* in the early Miocene of East Africa (Harrison 2010b). The divergence of lorisiforms is unlikely to have occurred before the first appearances of *Karanisia* at BQ-2 in Egypt (Seiffert, et al. 2003). We use 38 Ma as a conservative soft maximum.

## Platyrrhini | Pitheciidae-Callitrichidae 115.7 Ma – 33 Ma | SAB

The minimum age for the divergence of crown Platyrrhini is based on the first occurrence of the crown pitheciine *Proteropithecia* dated at 15.7 Ma (Kay, et al. 1998, Fleagle and Tejedor 2002). The minimum age could be as much 18 Ma if either (or both) *Soriacebus* or *Carlocebus* represent crown pitheciins (Fleagle, et al. 1987, Fleagle 1990, Bown and Fleagle 1993, Fleagle, et al. 1995, Rosenberger 2011). All of these taxa are known from the early and middle Miocene of Argentina (Fleagle and Tejedor 2002). The divergence of platyrrhines is unlikely to have occurred before the appearance of the crown catarrhine *Catopithecus* (33 Ma, Fayum, Egypt) although a recently published report has claimed a 36 Ma date for a stem platyrrhine from Peru (Bond, et al. 2015). It remains unclear how this older date was derived, however, and requires further substantiation.

## Atelidae | *Ateles-Alouatta* | 12.8 Ma – 18 Ma | SAB

The minimum age for the divergence of crown Atelidae (as recognized by Rosenberger, 2011; subfamily Atelinae of others) is based on the first appearance of the crown atelid *Stirtonia* (Hershkovitz 1970, Rosenberger 2011) at 12.8 Ma. Fossil atelids are known from the middle Miocene of Colombia and the Quaternary of Brazil and the Greater Antilles (MacPhee and Horovitz 2002). The divergence of atelids is unlikely to have occurred before the first appearance of the potential stem or crown atelid *Soriacebus* at 18 Ma (Bown and Fleagle 1993, Fleagle, et al. 1995).

## Cebidae | *Cebus-Saimiri* | 12.8 Ma – 18 Ma | SAB

The minimum age for the divergence of crown Cebidae is based on the first appearance of the crown cebid *Neosaimiri* (Stirton 1951, Hartwig and Meldrum 2002) dated at 12.8 from La Venta, Colombia. There are several older potential crown cebids including *Dolichocebus* and *Tremacebus* from Argentina and *Chilecebus* from Chile, all dated to around 20 Ma but it remains unclear how these taxa relate to the crown group. Recently, an additional potential crown cebid has been described from Panama (Bloch et al. 2016) dated at 20.9 Ma, which if substantiated would push the potential maximum bound to at least 21 Ma. Here we have accepted the notion that the divergence of cebids is unlikely to have occurred before the first appearance of the potential stem or crown atelid *Soriacebus* at 18 Ma (Bown and Fleagle 1993, Fleagle, et al. 1995) but acknowledge that this date could be as old as 21-22 Ma.

## Cercopithecidae | *Papionini-Cercopithecini* | 5 Ma – 23 Ma | SAB

The minimum age for the divergence of crown Cercopithecidae is based on the first appearance of *Parapapio* in the late Miocene (5 Ma) at Lothagam, Kenya. It is potentially possible that some specimens of *Parapapio* from Lothagam could be as old as 7.4 Ma (Jablonski and Frost 2010). The divergence of cercopithecids is unlikely to have occurred before the first appearance of the stem cercopithecoid *Prohylobates*. The oldest documented *Prohylobates* specimens are from Wadi Moghra in the Qatarra Depression, Egypt dated to approximately 19.5 Ma. However, we view *Kamoyapithecus*, dated at 25 Ma as a crown hominoid, which implies that at least stem cercopithecoids were in existence at that time. Given the controversial nature of our views on *Kamoyapithecus*, we have used a date of 23 Ma as the likely maximum divergence time for crown cercopithecids.

## Colobinae | Colobini-Presbytini | 9.8 Ma – 23 Ma | SAB

The minimum age for the divergence of crown Colobinae is based on the first appearance of *Microcolobus* in the middle Miocene (9.8 Ma) at Ngeringerowa, Kenya (Benefit and Pickford 1986, Jablonski and Frost 2010). The divergence of colobines is unlikely to have occurred before the first appearance of the stem cercopithecoid *Prohylobates*. The oldest documented *Prohylobates* specimens are from Wadi Moghra in the Qatarra Depression, Egypt dated to approximately 19.5 Ma. However, we view *Kamoyapithecus*, dated at 25 Ma as a crown hominoid, which implies that at least stem cercopithecoids were in existence at that time. Given the controversial nature of our views on *Kamoyapithecus*, we have used a date of 23 Ma as the likely maximum divergence time for crown colobines.

## APPENDIX 2

Here we briefly describe our new implementation of Bayes factor calculation in MCMCTree. Our approach is the thermodynamic integration-Gaussian quadrature method implemented recently in the program BPP. The mathematical details are given in Rannala and Yang (2017).

In this paper, we compare different rate models, *m*_*i*_, which differ only in the density of the branch rates while the prior on divergence times and the sequence likelihood are the same between the models. The posterior distribution of times (**t**) and rates (**r**) given the sequence data *D*, and given a clock model *m*_*i*_ is thus

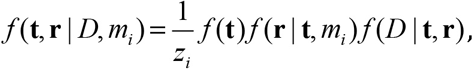

where

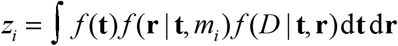

is the marginal likelihood of the data for model *m*_*i*_. Let *m_r_* be the model with highest marginal likelihood, and let BF_*ir*_ = *z*_*i*_/*z*_*r*_ be the Bayes factor of model *i* over model *r*. Then the posterior probability of model *i* is

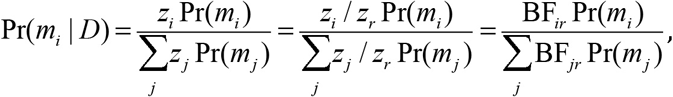

where the sum is over all models being tested, and Pr(mi) is the prior model probability.

We calculate *z_i_*, by sampling from the power posterior

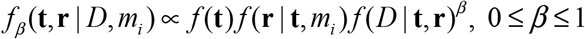

for a given *β* value. We choose *β*_*j*_ values to integrate between 0 and 1 according to the Gauss-Legendre quadrature rule. The estimate of log *z*_*i*_ is then given by the quadrature formula

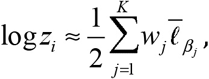

where *w*_*j*_ are the Gauss-Legendre quadrature weights, and 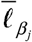 is the average of log-likelihood values, 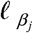, sampled from the power posterior with *β*_*j*_. The standard error of the estimate is given by

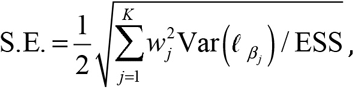

where the variance is calculated over the MCMC sample of 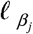, and ESS is the effective-sample size of 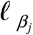.

